# Genomes of historical specimens reveal multiple invasions of LTR retrotransposons in *Drosophila melanogaster* populations during the 19^*th*^ century

**DOI:** 10.1101/2023.06.06.543830

**Authors:** Almorò Scarpa, Riccardo Pianezza, Filip Wierzbicki, Robert Kofler

## Abstract

Transposable element invasions have a profound impact on the evolution of genomes and phenotype. It is thus an important open question on how often such TE invasions occur. Based on strains sampled at different times, previous work showed that four different TE families invaded *D. melanogaster* during the 20^*th*^ century. Here we utilize the genomes of historical specimens to extend this work by another 100 years. We found that the LTR retrotransposons Blood, Opus and 412 spread in *D. melanogaster* in the 19^*th*^ century. These invasions were found to constitute second waves, as degraded fragments were found for all three TEs. We show that two of the three invasions, Opus and 412, led to pronounced geographic heterogeneity, likely due to founder effects during the invasions. Finally, we identified species from the simulans complex as the likely origin of the TEs. In total, seven TE families invaded *D. melanogaster* during the last 200 years, increasing the genome size by 1.2 Mbp. Based on the analysis of strains and specimens sampled at different times, we provide a detailed timeline of TE invasions, making *D. melanogaster* the first organism where we could infer the invasion history of TEs during the last two centuries.

## Introduction

Transposable elements (TEs) are short stretches of DNA that selfishly multiply within genomes. These elements can be broadly classified into two main classes: Class I, also known as retrotransposons, and Class II, referred to as DNA transposons. Retrotransposons propagate by a copy-and-paste mechanism that involves an RNA intermediate as a template, whereas DNA transposons directly relocate to new genomic locations using a cut-and-paste mechanism [21, 70, 7]. For retrotransposons usually long-terminal repeat (LTR) and non-LTR TEs can be distinguished [21, 29, 70]. TEs are highly successful, having invaded virtually all eukaryotic species investigated so far [70].

Since many TE insertions are likely deleterious [17, 49], host organisms evolved elaborate defense mechanisms against them [9, 44, 56]. In *D. melanogaster* the defense against TEs is based on piRNAs (PIWI-interacting RNAs), i.e. small RNAs with a size between 23–29nt, that repress TE activity at the transcriptional and post-transcriptional level [9, 25, 60, 39]. These piRNAs are largely derived from discrete genomic loci, the piRNA clusters. [9]. In *D. melanogaster* piRNA clusters account for about 3% of the genome [9]. It is thought that a TE invasion is stopped when a copy of the invading TE jumps into a piRNA cluster, which triggers the emergence of piRNAs complementary to the invading TE [76, 43, 23, 75, 48]. One particularly important component of the piRNA pathway is the ping-pong cycle which amplifies the amount of piRNAs by alternately cleaving sense and antisense transcripts of TEs [9, 25]. Activation of the ping-pong cycle may be necessary for silencing an invading TE completely [58]. Once a TE is inactivated all insertions of a family will decay by accumulating mutations over time. Eventually the TE might not be able to mobilize anymore, resulting in the death of a TE family [4]. One strategy for escaping this inactivation, thereby ensuring the long-term persistence of a TE is horizontal transfer (HT) to a naive species not having the TE [4]. Such HT may trigger TE invasions in the naive species, that are than in turn silenced by the host defence [4]. HT is probably abundant. For example, a study investigating 195 insect species identified about 2000 HTs of Tes [50]. Several HT of TEs were also reported in *D. melanogaster* [2, 55]. In agreement with this, most LTR families in *D. melanogaster* are likely of recent origin, possibly as young as 16.000 years [3, 8]. Moreover, four different TEs invaded *D. melanogaster* populations during the last 100 years [57]. Three of these TE invasions - the P-element, Hobo, and the I-element - were discovered due to phenotypic effects caused by the activity of the TE, i.e. the hybrid dysgenesis (HD) symptoms [33, 32, 1, 51, 13, 14, 11, 6]. Crosses between males having a TE and females not having it frequently lead to diverse phenotpyic effects, such as atrophied ovaries, whereas no phenotypic effects can be found in reciprocal crosses [33]. By sequencing some of the oldest available *Drosophila* strains, we recently showed that a fourth TE, Tirant, also invaded *D. melanogaster* during the last 100 years [57]. We did not notice any HD symptoms caused by Tirant, which may account for the late discovery of the Tirant invasion [57]. Hobo, I-element, and Tirant likely spread in *D. melanogaster* in multiple waves as degraded and fragmented copies of these TEs could be found in all investigated strains [57]. Solely the P-element did not show similarity to any sequence of the *D. melanogaster* genome. Interestingly, we found that the Tirant composition showed geographic heterogeneity where populations from Tasmania carried slightly different Tirant variants than other populations, likely due to a founder effect during the invasion [57]. In agreement with such a geographic heterogeneity of the TE composition a recent work identified diverse TE lineages (i.e. SNPs showing correlated allele frequencies across different samples) for multiple TE families in *D. melanogaster* [54]. By investigating the presence/absence of TE families in strains sampled at different times we were able to reconstruct the invasion history of *D. melanogaster* during the last 100 years [57]. Tirant invaded natural *D. melanogaster* populations first (1930-1950), followed by the I-element, Hobo and lastly by the P-element [57]. The oldest of these strains used for reconstructing the invasion history of *D. melanogaster*, Oregon-R and Canton-S, were sampled between 1925 and 1935 [57]. Recently the genomes of 25 historical *D. melanogaster* specimens became publicly available which provides us with an opportunity to extend the invasions history of *D. melanogaster* by another 100 years [59]. Six strains were sampled around 1800 in Lund (Sweden; early 1800), two around 1850 in Passau (Germany; mid 1800), one around 1900 in Lund (late 1800), and 16 around 1933 in Lund [59]. By analysing the genomes of these historical specimens we found that the three LTR transposons - Blood, Opus, 412 - invaded *D. melanogaster* populations likely between 1850 and 1933. All three TEs are LTR retrotransposons, where Blood and 412 do not have an envelope protein and belong to the Gypsy/mdg1 superfamily whereas Opus (also known as Nomad) has an envelope protein and belongs to the Gypsy/Gypsy superfamily [29]. Similarly to Tirant (Gypsy/Gypsy superfamily), Opus may thus form virus-like-particles that could infect the germline. All three TEs have a similar size (Blood=7, 410bp, 412=7, 567bp, Opus=7, 512bp), between 2-4 annotated ORFs (Blood=3, 412=4, Opus=2) and LTRs with a similar size (Blood=398bp, 412=514bp, Opus=518bp)[52]. We also found short degraded fragments (20-30% divergence) for Opus, 412 and Blood in all investigated specimens, which likely represent remnants of an ancient wave of an invasion. By investigating TE specific SNPs in extant populations we found that the composition of Opus and 412 - but not of Blood - varies among populations, where especially populations from Zimbabwe carry slightly different variants than other populations. This geographic heterogeneity could be due to founder effects during the invasion of Opus and 412. We suggest that HT from a species of the simulans-complex likely triggered the invasions of Blood, Opus and 412. By jointly analysing the genomes of strains and specimens sampled at different times we were able infer a invasion history of TEs in *D. melanogaster* during the last 200 years: Blood, Opus and 412 spread between 1850-1933, followed by Tirant and the I-element between 1933-1950, Hobo around 1950, and finally the P-element between 1960-1980 (see also [57]). To our knowledge, this makes *D. melanogaster* the first species where it is feasible to infer a detailed invasion history of TEs during the last two centuries.

## Results

### The LTR transposons Blood, 412 and Opus likely invaded dmel between 1800 an 1933

Sequencing of the oldest available *D. melanogaster* strains, sampled between 1925 and 1938, revealed invasions of four different TEs (Tirant, Hobo, I-element, P-element) in natural populations during the last 100 years [57]. The publication of the genomes of 25 historical *D. melanogaster* specimens, collected between 1800 and 1933, provides us with the opportunity to investigate whether additional TE invasion occurred between 1800 and 1933 [59]. To do this, we compared the abundance of TEs in the historical specimens to more recently collected strains. We downloaded the publicly available reads, filtered or trimmed them to a size of 100bp, aligned the reads to the consensus sequences of TEs in *D. melanogaster* [52] and estimated the abundance of TEs with our tool DeviaTE [69] (for an overview of the data used in this study see supplementary table S1). For each TE family, DeviaTE normalizes the average coverage of a TE (e.g. 121) to the average coverage of single copy genes (e.g. 12), which allows inferring the TE copy number per haploid genome (e.g. 10.1 = 121*/*12). We first compared the TE abundance between a strain collected around 1800 (18SL13) and the strain Harwich. Harwich was collected around 1967 and should thus contain copies of all TEs that invaded *D. melanogaster* populations during the last 100 years (i.e. Tirant, Hobo, I-element, P-element; [57]). As expected we found a strong overrepresentation of Tirant, Hobo, the I-element and the P-element in Harwich (fig. 1A blue). Surprisingly, we additionally found that 412, Blood and Opus are highly overrepresented in Harwich as compared to 182L13 (fig. 1A red). A comparison between 182L13 and a strain collected around 1938 (Lausanne-S) showed an overrepresentation of 412, Blood and Opus in Lausanne-S but not of Tirant, the I-element, Hobo, and the P-element (fig. 1A red). By contrast, a comparison between Lausanne-S and Harwich solely revealed an overrepresentation of Tirant, the I-element, Hobo and the P-element but not of Opus, Blood and 412 (supplementary fig. S1).

**Figure 1:**
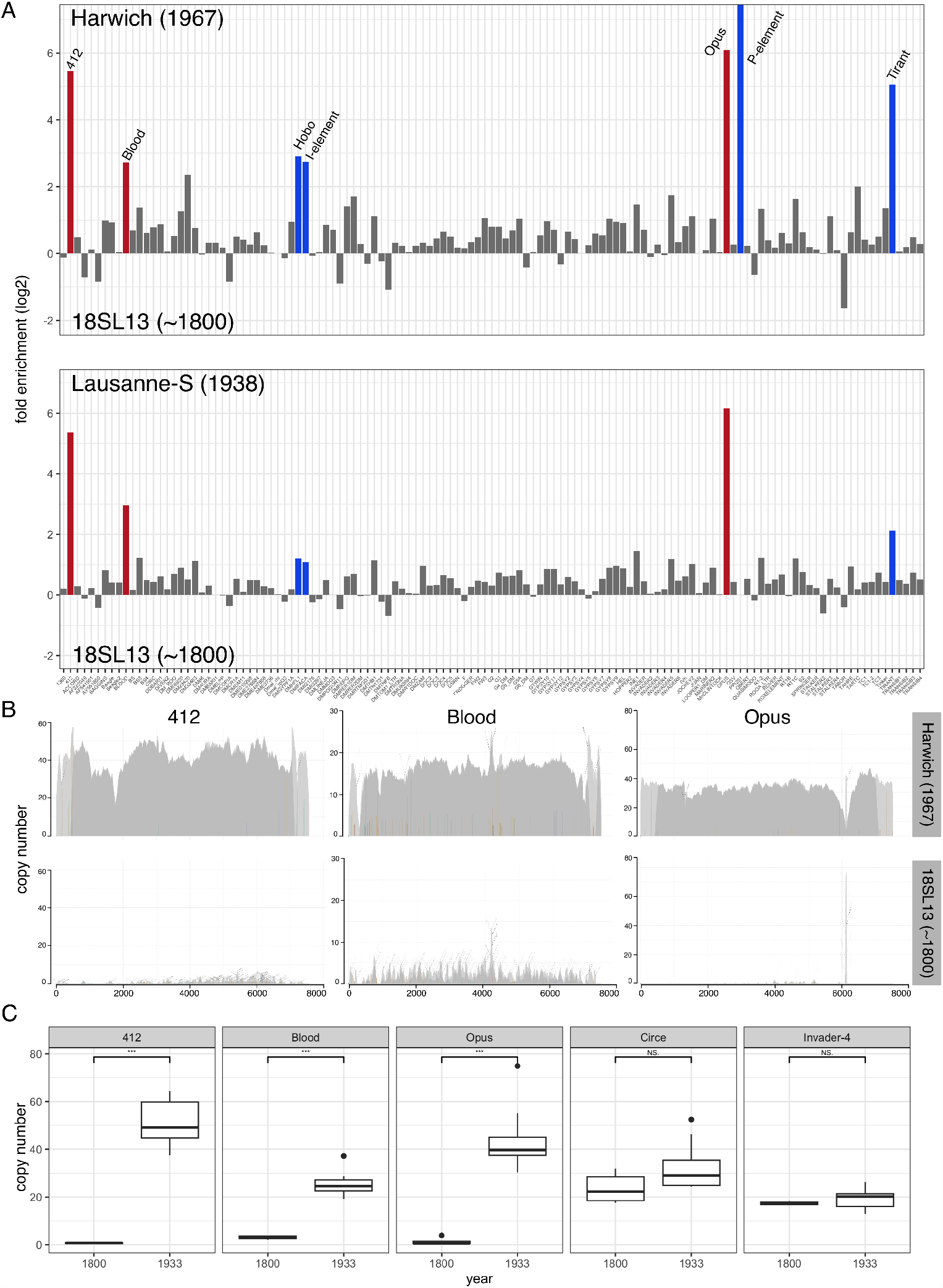
Genomes of historical *D. melanogaster* specimens suggest that the LTR retrotransposons Opus, Blood and 412 invaded *D. melanogaster* populations in the 19^*th*^ century. A) Differences in TE abundance between a strain collected around 1800 (182L13) and strains collected in 1967 (Harwich) or 1938 (Lausanne-S). For each TE family (x-axis) we show the difference in TE copy numbers per haploid genome as estimated with DeviaTE (y-axis). Note that Blood, Opus and 412 (red bars) show a strong enrichment between strains collected at 1800 and 1938. As expected, the previously described invasions of Hobo, I-element, P-element and Tirant (blue bars) are only revealed in the comparison with the more recently collected strain Harwich. B) Abundance and diversity of 412, Blood and Opus in a strain collected around 1800 and 1938. The short reads were aligned to the consensus sequences of these TEs and visualized with DeviaTE. The coverage of the TEs was normalized to the coverage of single copy genes. SNPs and indels are shown as colored lines. Coverage based on unambiguously and ambiguously aligned reads is shown in dark and light grey, respectively. C) 4 Copy numbers of 412, Opus, Blood in historical specimens collected around 1800 (9 samples) and 1933 (16 samples). As controls Circe and Invader-4 are included. The significance was computed with Wilcoxon rank sum tests.

An analysis of the coverage revealed that 412, Blood and Opus have a uniformly elevated coverage in Harwich as compared to 182L13 (fig. 1B), which suggests that overrepresentation of these three TEs in Harwich is not an alignment artefact (e.g. due to low complexity regions). Such an uniformly elevated coverage in specimens sampled around 1933 as compared to those sampled around 1800 can be found for all of the analysed samples (supplementary figs S2, S3, S4). Comparing the sequences of these three TEs with BLAST did not reveal any sequence similarity, ruling out crossmapping among these three TEs. Solely a few high frequency SNPs can be found for the three TEs in Harwich, which suggests that most of the reads align without mismatch to the consensus sequence to the TEs (fig. 1B). By contrast only few highly diverged reads align to Opus, 412 and Blood in 182L13 (fig. 1B). The estimated copy numbers of 412, Opus and Blood are significantly higher in specimens collected around 1800 as compared to specimens collected at 1933, whereas no significant differences could be found for other TEs such as Circe and Invader4 (fig. 1C). An analysis independent of DeviaTE, solely based on the number of reads aligning to the TEs confirms this significant difference in the abundance of 412, Blood and Opus when comparing strains sampled in 1800 and 1933 (supplementary fig. S5). So far we solely considered reads with a length of at least 100bp. However most of the reads from historical samples are degraded with a length of 50bp [59]. We thus repeated these analysis with reads of 50bp (longer reads were trimmed) and again found significantly elevated copy numbers for Blood, Opus and 412 but not for Circe and Invader-4 between strains collected around 1800 and 1933 (supplementary fig. S6). Our data thus suggest that 412, Blood and Opus invaded natural *D. melanogaster* populations between 1800 and 1933. To further test this hypothesis we investigated the length and divergence of these TEs in four high-quality assemblies (mostly based on long reads) of *D. melanogaster* (Canton-S, Iso1, Pi2, Dgrp-732 [27, 18, 74]). For recently active TEs we expect to find multiple full-length insertions with a high similarity to the consensus sequence. Indeed, in each analysed strain we found multiple full-length insertions of Blood, Opus and 412 that showed little divergence to the consensus sequence (*<* 1%; supplementary fig. S7). However, for each of these TEs we additionally found several fragmented and highly diverged (20-30%) insertions (supplementary fig. S7). These diverged insertions could account for the few reads mapping to these three TEs in historical samples collected around 1800 (fig. 1B). The degraded copies of Blood, Opus and 412 are likely the remnants of ancient invasions of these TEs. Similarly to Tirant, Hobo and the I-element (but not the P-element) the three TEs (Opus, Blood and 412) likely invaded *D. melanogaster* in multiple waves.

In summary we suggest that the LTR retrotransposons Blood, Opus, and 412 invaded natural *D. melanogaster* populations in the 19^*th*^ century. These recent invasions likely constitute second waves of invasions as we found highly degraded fragments of these TEs in all investigated strains.

### The invasion history of TEs in *D. melanogaster* during the last 200 years

In a previous work we inferred the invasion history of TEs in natural *D. melanogaster* populations by sequencing different laboratory strains collected during the last century [57]. The oldest available strains, Oregon-R and Canton-S, were collected around 1925 to 1936. Given the availability of the museum specimens we aim to extend this work by another 100 years, thus inferring the invasion history of TEs in *D. melanogaster* during the last 200 years (until 1800). We estimated the copy numbers of the seven TEs that recently invaded *D. melanogaster* (Blood, Opus, 412, Tirant, the I-element, hobo, the P-element) in the historical specimens as well as in diverse strains sampled during the last century (for an overview of all investigated strains and specimens see supplementary table S1). We trimmed reads to a size of 100bp, mapped them to the consensus sequences of TEs in *D. melanogaster* and estimated the copy numbers with DeviaTE (see above; [69]). Opus, Blood and 412 were largely absent in all strains sampled until ≈1850 (fig. 2A; supplementary table S1). We noticed a sudden increase in the number of reads mapping to Opus, Blood and 412 starting in some samples collected in the late 1800, where these three TEs were present in all specimens collected after 1933 (fig 2A; supplementary table S1). We thus suggest that Opus, Blood and 412 invaded natural *D. melanogaster* populations between 1850 and 1933 (fig 2B; supplementary table S1). To provide the complete invasion history of TEs during the last 200 years we also estimated the abundance of the TE families analyzed in our previous work, i.e. Tirant, Hobo, I-element and the P-element [57]. In agreement with our previous work our data suggest that Tirant invaded *D. melanogaster* populations between 1933-1950, followed by the I-element, Hobo and lastly by the P-element (fig 2B; supplementary table S1[57]). In summary we suggest that the LTR retrotransposons Opus, Blood, 412 invaded natural *D. melanogaster* populations between ≈1850 and 1933, Tirant and the I-element between 1933 and 1950, Hobo around 1950 and the P-element between 1960-1980. To our knowledge *D. melanogaster* is the first species where the history of TE invasions during the last centuries could be inferred.

**Figure 2:**
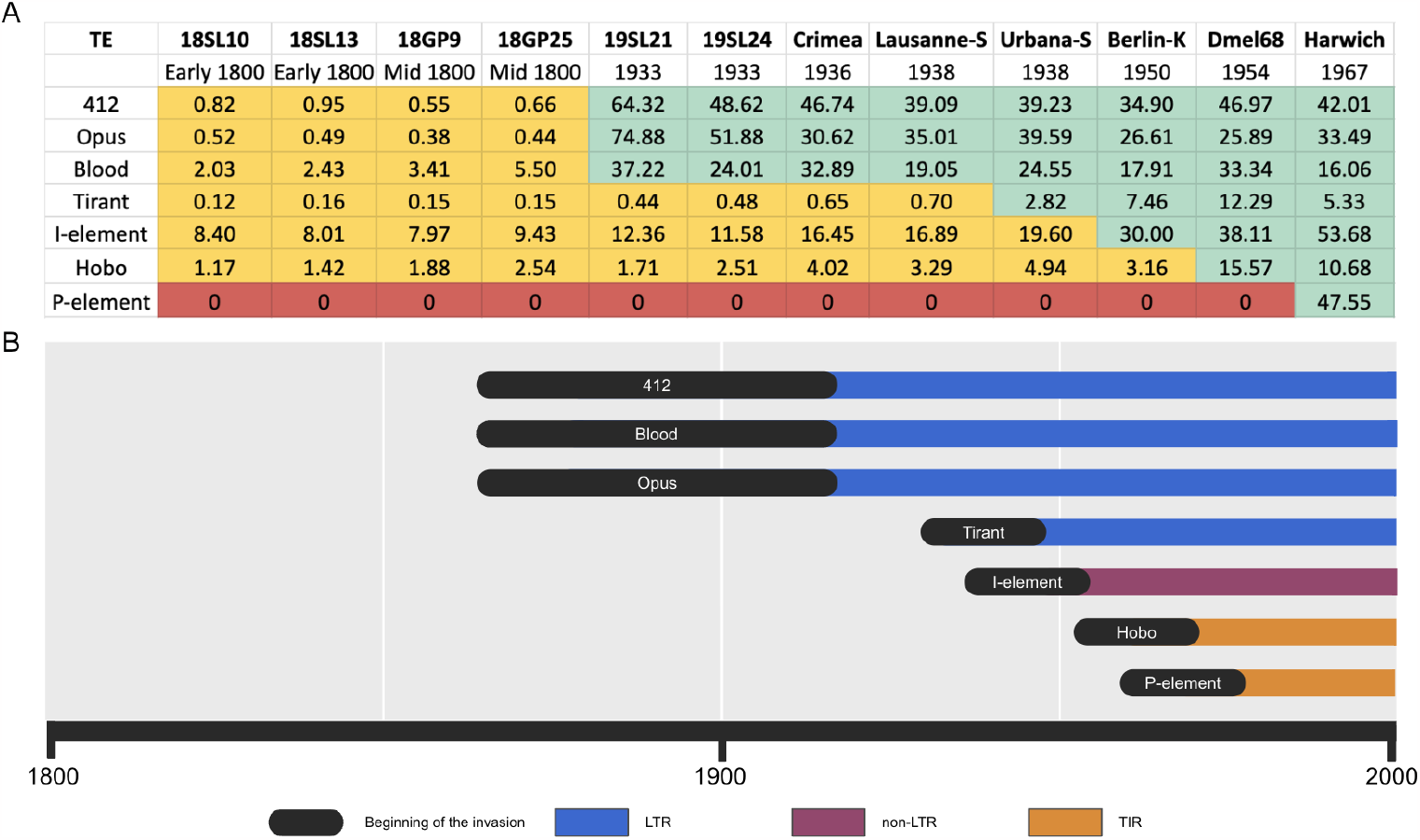
History of TE invasions in *D. melanogaster* during the last 200 years. A) Overview of the abundance of Blood, Opus, 412, Tirant, the I-element, Hobo and the P-element in strains or specimens sampled at different years. For each TE family we classified the abundance into the categories: absence of TE (red), solely degraded copies are present (likely remnants of ancient invasions; yellow), presence of non-degraded copies with a high similarity to the consensus sequence (green). The numbers represent copy numbers in haploid genomes as estimated by DeviaTE. B) Timeline of TE invasions in *D. melanogaster*. The width of the black bars indicates the range of uncertainty of the invasions.

### Blood, Opus and 412 are silenced by the piRNA pathway in natural populations

We next asked whether Blood, Opus and 412 are under host control by the piRNA pathway in extant populations. To address this question we interrogated small RNA data from the Global Diversity Lines (GDL) which comprise 85 *D. melanogaster* strains sampled after 1988 from five different continents (Africa - Zimbabwe, Asia - Beijing, Australia - Tasmania, Europe - Netherlands, America - Ithaca [24]. The small RNA were sequenced for 10 out of the 85 GDL strains, where two strains were selected from each continent [42]. We found abundant sense and antisense piRNAs distributed over the entire sequence of the three TEs in all 10 GDL strains (supplementary fig. S8). In the germline, the amount of piRNAs complementary to a TE is greatly amplified by the ping-pong cycle [9, 25]. Activity of this ping-pong cycle is likely necessary to establish host control over an invading TE [10, 58]. An active ping-pong cycle generates a characteristic overlap between the 5’ positions of sense and antisense piRNAs, i.e. the ping-pong signature [9, 25]. All three TEs show noticeable ping-pong signatures in the 10 analysed GDL strains (supplementary fig. S9). We thus argue that Blood, Opus and 412 are controlled by the piRNA pathway in extant *D. melanogaster* populations.

### The composition of Opus and 412 but not of Blood varies among extant populations

We previously found that the Tirant composition varies among populations where especially populations from Tasmania carried slightly different variants than populations from other geographic locations [57]. To investigate whether a geographic heterogeneous compositions can also be found for Blood, Opus and 412 we investigated the composition of these TEs in the 85 GDL strains ([24]; for an overview of all analysed strains see supplementary table S2). For each TE family we identified SNPs and estimated the allele frequencies of the SNPs. Notably, in this work a SNP refers to a variant among dispersed TE copies. Our allele frequency estimates thus reflect the TE composition within a particular strain (e.g. if 15 Blood insertions in a strain carry a ‘G’ at a particular site and 5 an ‘A’, the frequency of G at this site is 0.75). We used UMAPs to summarize differences in the TE composition among the GDL strains. UMAPs are widely used in population genetics [16]. In contrast to principal component analysis (PCA), UMAPs have the advantage of summarizing information from all axis of variation in a sample (not just the main axis), thereby revealing fine-scale patterns in population structure [15, 16]. The main disadvantage of UMAPs is that the distance among samples in the plot does not reflect the genetic distance [16]. We first confirmed that UMAPs capture the previously reported geographic heterogeneity of Tirant (fig. 3). We found that Opus and 412 but not Blood show geographic heterogeneous compositions. For Opus populations from Tasmania and Zimbabwe are distinct, while for 412 populations from Zimbabwe and to a minor extend from Bejing form distinct clusters. To rule out that this geographic pattern is merely due to the ancient fragments of these TEs we repeated these UMAPs by excluding all sites having a coverage in specimens collected around 1800 but found the same clusters (supplementary fig. S10). These clusters can also be discerned with PCA (using the first two principal components) but less clearly than with the UMAPs (supplementary fig. S11). We next investigated the reasons for these distinct clusters in the UMAPs. Therefore, we aimed to identify diagnostic SNPs for these TEs, i.e. SNPs that are abundant in a population of interest but rare in all other populations (supplementary fig. S12). We found several diagnostic SNPs with a high-frequency for Tirant in Tasmania, Opus in Zimbabwe and Tasmania and 412 in Zimbabwe and Bejing (supplementary fig. S12). No diagnostic SNPs with a high-frequency were found for Blood (supplementary fig. S12). We thus argue that the diagnostic SNPs reflect the clusters of the UMAP. For an overview of the most abundant diagnostic SNPs see supplementary table S3. Differences in the TE composition among the GDL populations are thus likely responsible for the geographic heterogeneity observed for Tirant, Opus and 412. Interestingly the geographic pattern seen for 412 resembles the pattern found with “neutral” autosomal SNPs, where populations from Zimbabwe and Bejing also form distinct clusters from the other populations [24]. The geographic heterogeneity in the TE composition could be either due to founder effects during the TE invasions or to demographic processes (see Discussion).

**Figure 3:**
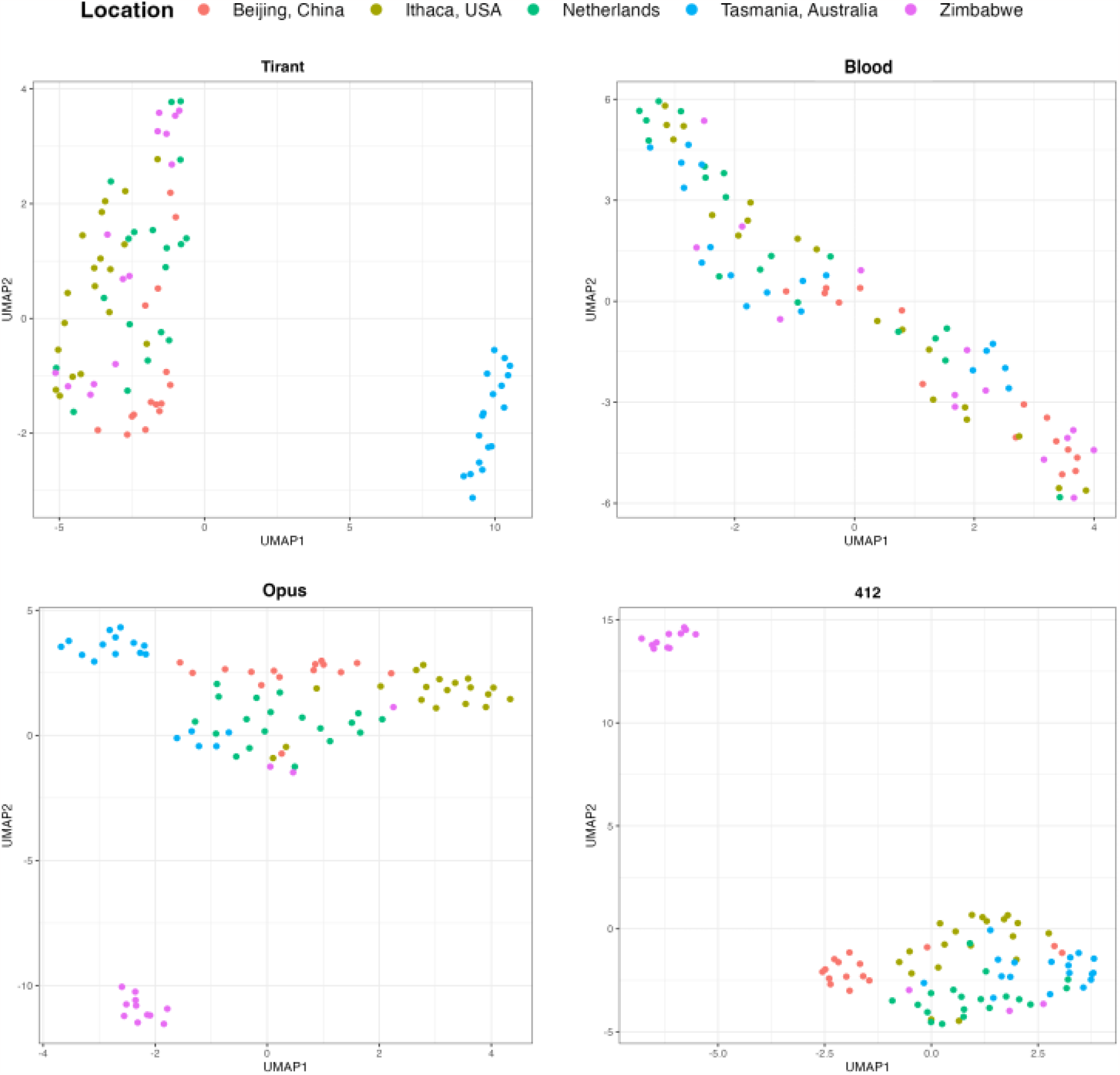
UMAPs for Tirant, Blood, Opus and 412 in the 85 GDL strains. UMAPs are based on the allele frequency of TE specific SNPs. Note that Tirant, Opus and 412 show population structure, likely due to founder effects during the invasion.

### Origin of horizontal transfer

Here we propose that Blood, Opus and 412 recently invaded *D. melanogaster* populations, likely following horizontal transfer (HT) from a different species. To identify the possible source of the HT we investigated the genomes of 101 long-read assemblies of different drosophilid species [34]. We also included the long-read assemblies of recently collected *D. melanogaster* and *D. simulans* strains (Pi2, SZ232 [73, 72]). We reasoned that the species that acted as donor for HT should have insertions with a high similarity to the consensus sequence of the TEs in *D. melanogaster*. Using RepeatMasker we identified TE insertions in these 103 assemblies and estimated the similarity of the insertions to the consensus sequence of *D. melanogaster*. We first tested if this approach allows us to reproduce the likely donor species for Tirant, I-element, P-element, and Hobo. Apart from *D. simulans*, which recently acquired the P-element [37, 26], we find that *D. willistoni* carries P-element insertions that are most similar to the P-element in *D. melanogaster* (fig. 4). This is in agreement with previous work suggesting that a species from the willistoni or saltans group is the likely source of the P-element in *D. melanogaster* [14]. For Hobo, the I-element, and Tirant, a species from the *D. simulans* complex was suggested as the likely donor [57, 13, 61, 41]. In agreement with this we also find that species from the *D. simulans* complex have insertions that are most similar to the consensus sequence of Tirant, Hobo and the I-element (fig. 4). We further found that species from the *D. simulans* complex have insertions that are most similar to the consensus sequence of Blood, Opus and 412 (fig. 4). We thus suggest that a species from the *D. simulans* complex is the most likely source of the HT that triggered the invasions of Blood, Opus and 412 in natural populations of *D. melanogaster*.

**Figure 4:**
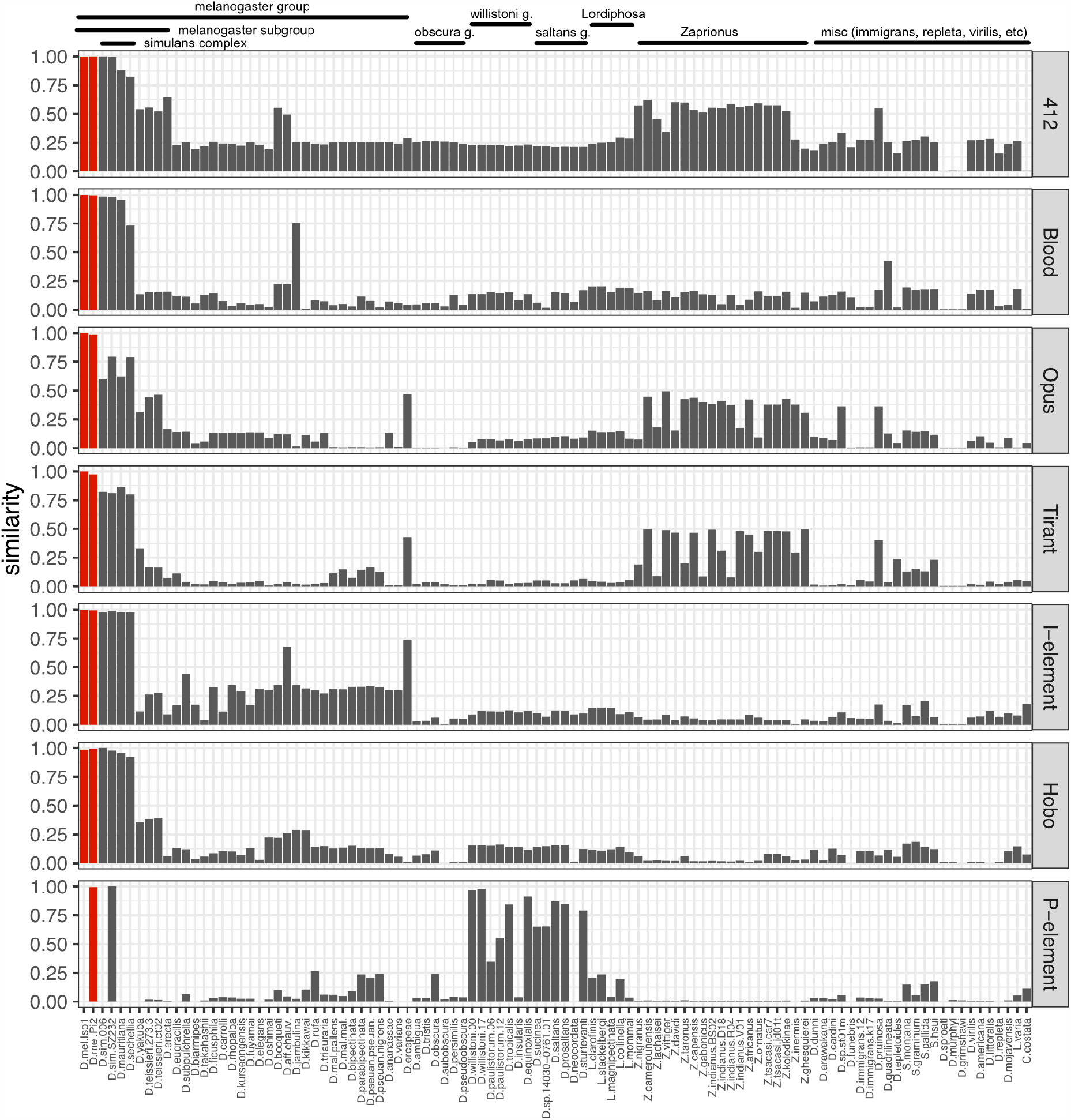
Possible origins of the seven TE families that invaded *D. melanogaster* populations during the last 200 years. Data are shown for 103 long-read assemblies of diverse drosophilid species (red *D. melanogaster*). The barplots show the similarity of TE insertions in a given assembly to the consensus sequence of the TE family. For example, a value of 0.9 indicates that at least one TE insertion in the assembly has a high similarity (≈ 90%) to the consensus sequence of the TE. Apart from the P-element, which was likely transmitted from *D. willistoni* to *D. melanogaster*, all other TE families have insertions with the highest similarity in species from the *D. simulans* complex. HT of insertions from the *D. simulans* complex thus likely triggered the invasions of Blood, Opus and 412 in *D. melanogaster* populations.

## Discussion

Taking advantage of publicly available genomes from historical *D. melanogaster* specimens we showed that the LTR retrotransposons Blood, Opus and 412 invaded natural *D. melanogaster* populations between ≈1850 and 1933. Solely a few degraded reads aligned to these TEs in any specimen collected until ≈1850, but a substantial number of reads with a high similarity to the consensus sequence aligned to these TEs in all specimens collected in 1933 (fig. 1). This finding is robust to different methods for estimating TE copy numbers and to different read length (supplementary figs. S5,S6). The presence of multiple full-length insertions with a high similarity to the consensus sequence in multiple long-read assemblies of different *D. melanogaster* strains also supports the idea that Blood, Opus and 412 were recently active (supplementary fig. S7). A high similarity among insertions of Blood, Opus and 412 was also noticed in previous works (Opus = Nomad) [8, 3]. Insertions of Blood, Opus and 412 are largely segregating at a low frequency in natural *D. melanogaster* populations, which further suggests that these TEs were recently active [35, 38].

By analysing strains and specimens collected at different times we provide an updated history of TE invasions *D. melanogaster* populations during the last 200 years. Our data suggest that seven TEs invaded natural *D. melanogaster* populations during the last 200 years. The four oldest invasions - Blood, Opus, 412 and Tirant which invaded between 1850 and 1950 - were due to LTR retrotransposons (fig. 2). Our findings thus provide strong support for the work of Bergman et al. [3] who, based on an analysis of substitutions among TE sequences, suggested that most LTR retrotransposons in *D. melanogaster* are of very recent origin (*<* 100, 000 years, possibly even *<* 16, 000 years). We found that species from the *D. simulans* complex carry insertions that are very similar to the consensus sequences of Blood, Opus and 412 in *D. melanogaster* (fig. 4). Therefore, we propose that HT from a species of the *D. simulans* complex triggered the invasion of Blood, Opus and 412 in *D. melanogaster*. One problem with this analysis is that we cannot infer the direction of the HT. It is for example possible that a HT from *D. melanogaster* triggered invasions of Blood, Opus and 412 in a species of the *D. simulans* complex and not vice versa. Therefore it is feasible that the invasions of 412, Opus, and Blood were not triggered by HT from a drosophilid species but rather from an entirely different species. For example, mariner is a highly successful TE that spread in many diverse drosophilid and non-drosophilid species [53]. However, an analysis of 99 long-read assemblies of different insect species did not identify any insertions similar to Blood, Opus and 412 (supplementary fig. S13; supplementary table S5).

A possible explanation for the high number of TE invasions triggered by HT from a species of the *D. simulans* complex was suggested by [4]: TEs may be frequently horizontally transmitted back-and-fourth between related species. As a consequence TEs from related species may periodically reinfecting each other, thus ensuring the long term persistence of the TEs [4]. Our finding that HT from a species of the *D. simulans* complex likely triggered the invasion of 6 out of 7 TEs that invaded *D. melanogaster* during the last 200 years supports this hypothesis. The presence of degraded insertions in addition to full-length insertions for 6 out of these 7 TEs (supplementary fig S7) further suggests that these TEs invaded *D. melanogaster* in multiple waves, again in agreement with the hypothesis of Blumenstiel [4].

Based on an analysis of ancient genomes of Drosophila we estimate that seven TEs invaded *D. melanogaster* populations during the last 200 years. Although HT is occurring frequently among insect species [50], our work raises the important open question of whether such a high rate of TE invasions (one invasion all ≈30 years) can also be found in other species. It will be important to test this hypothesis with additional species. Although strains sampled at different time points will only be available for few species, an analysis of historical museum specimens should in principle be feasible for many diverse species.

A related question is whether the observed high rate of TE invasions during the last 200 years is even representative for the evolution of *D. melanogaster*. The observed 200 years could represent an unusual accumulation of TE invasions. If we roughly interpolate the rate of invasions (one TE invasion all 30 years), the ≈121 TE families found in the genome of *D. melanogaster* could have been acquired by HT during the last 3600 years. However, given that we can still identify TEs that invaded the *D. melanogaster* genome 2-5 million years ago, such as Ine1, jockey2, Helena or Cr1a [3, 62], we would expect one invasion all 16,000 years (2 million years / 121 families) or one invasion all 8,000 years if two waves of invasions are assumed for each TE family. We thus think that the rate of TE invasions observed during the last 200 years in *D. melanogaster* is a unusually high. This raises the question of which events might have triggered such a high rate of HTs in the last 200 years. One possible explanation could be the recent habitat expansion of *D. melanogaster* into the Americas and Australia [68, 57]. *D. melanogaster* originated in tropical sub-Saharan regions of Africa, started to colonize the rest of the World about 10,000 years ago, spread from the middle East into Europe about 1800 years ago, and finally spread to the Americas and Australia about 100-200 years ago [31, 5**?**, 64]. Habitat expansion may bring species into contact that were previously isolated, thus generating novel opportunities for HT among species. An illustrative example is the P-element in *D. melanogaster*, which was likely acquired from *D. willistoni* after *D. melanogaster* entered the habitat of *D. willistoni* in South America [19]. The fact that the invasion of Blood, Opus and 412 spread around the same time that *D. melanogaster* spread into North America and Australia argues in favor of the habitat expansion. Given that Blood, Opus and 412 spread in *D. melanogaster* populations around the same time (1850-1933), we speculate whether these three invasions could have been triggered by a single event, such as an introgression from *D. simulans* into *D. melanogaster*. Our finding that the three TEs have insertions with high similarity in species of the *D. simulans* complex is consistent with a common origin of the three invasions. Further genomes of historical specimens collected between 1850 and 1933 could provide a more detailed resolution of this period, thus enabling us to estimate whether these three TEs spread at the same time.

Out of the four TEs that invaded *D. melanogaster* during the last century three TEs - the I-element (non-LTR), Hobo (DNA transposon), the P-element (DNA transposon) - cause diverse hybrid dysgenesis (HD) effects. Crosses among males having the TE with females not having the TE typically lead to offspring where the TE is active and this TE activity can lead to different phenotypic effects such as atrophied ovaries [33, 47]. This raises the question on whether Blood, Opus and 412 could also induce HD symptoms. Answering this question requires both, strains that have these TEs and strains that do not have them. Since the oldest available lines of *D. melanogaster* were collected around 1925-1933 we do not have any strains that are devoid of recent Blood, Opus and 412 insertions. The question could thus solely be answered by artificially introducing these TEs into naive strains (for example using a different species such as *D. erecta* [58]). Given that we did not detect any HD symptoms for Tirant, i.e. the sole LTR retrotransposon that invaded during the last 100 years [57], we suspect that the LTR retrotransposons Blood, Opus and 412 might also not induce any HD symptoms.

Based on SNPs found in the TEs we show that the composition of Opus and 412 - but not of Blood - varies among populations. For Opus specimens from Zimbabwe and Tasmania form distinct groups and for 412 specimens from Zimbabwe and Bejing (fig. **??**). A previous work based on PCAs additionally found that the composition of Tirant but not of the I-element, Hobo and the P-element varies among populations [57]. Here we confirm that the composition of Tirant varies among populations using UMAPs, where especially population from Tasmania carry distinct variants. With UMAPs we also confirm that the I-element, Hobo and the P-element do not show population structure (supplementary fig. S14). We think that two different processes could lead to a heterogeneous TE composition among extant populations, founder effects during the invasion and demographic processes. An analysis of “neutral” autosomal SNPs revealed that the populations from Zimbabwe and Bejing form distinct groups (based on the first two principal components [24]), which is very similar to the population structure that we observed for 412 (and to a more minor extent for Opus, where solely the population from Zimbabwe forms a distinct cluster). This raises the possibility that demographic processes shaped the composition of TEs in the extant population. There are however two problems with this hypothesis. First the geographic pattern varies among the TE families (e.g. Tasmania is a separate cluster for Tirant while Zimbabwe is a distinct cluster for Opus and 412) and several TE families show no discernible geographic pattern (Blood, I-element, Hobo and P-element). If demographic processes shaped the TE composition we expect that all TE families show the same geographic pattern. Second, Opus and 412 invaded *D. melanogaster* populations during 1850-1933. If demographic processes shaped the TE composition than they must have accomplish this during the last 150 years, which then raises the possibility that the geographic pattern seen with autosomal SNPs was also generated during these last 150 years. However the pattern seen for “neutral” autosomal SNPs are likely due to the out-of-Africa migration of *D. melanogaster* several thousand years ago [24] and not the result of recent demographic processes. We thus favor the hypothesis that founder effects during the invasions of the TEs are responsible for the observed geographic pattern seen for Tirant, Opus and 412. For example a few Opus insertions, with a slightly different composition, may have triggered the invasion of populations from Zimbabwe. As a result the populations from Zimbabwe will end up with a slightly different TE composition than other populations (similar to a founder effect when a new population is established). The geographic pattern observed for 412 (and to some extent for Opus) and the “neutral” autosomal SNPs might thus have emerged twice independently. The similarity of the pattern could just reflect the fact that invading TEs and migrating flies need to overcome the same barriers (e.g. the Sahara).

The seven TE invasions during the last 200 years had a substantial impact on the *D. melanogaster* genome. Due to these invasions the genome size of *D. melanogaster* increased by ≈1.2Mb in a short period of time (based on Pi2; supplementary table S4). These novel TE insertions could provide variation driving adaptation [22, 12], generate novel piRNA clusters (especially Blood and 412 frequently form ‘de novo’ clusters [65]), remodel gene regulatory networks [20] and generate diverse structural variants [30]. The high rate of TE invasions during the last 200 years may thus have had a substantial impact on the evolution of *D. melanogaster*.

## Materials and Methods

### Analysis of genomic DNA

We analysed the TE content in genomic DNA of *D. melanogaster* samples from three different publicly available data sets: the Global Diversity Lines [24] (accession number: PRJNA268111), lab strains collected at different times [57] (accession number: PRJNA634847) and the historical museum specimens [59] (accession number: PRJNA945389). For an overview of the analysed samples see supplementary tables S1, S2. We downloaded the files using wget, checked the md5 sum and trimmed the reads to 100bp. To investigate the robustness of our results we performed an additional analysis were all reads were trimmed to 50bp. The reads were mapped to a database consisting of the consensus sequences of TEs [52] and three single copy genes (*rhino, trafficjam, rpl32*) with bwa bwasw (version 0.7.17-r1188) [40]. Several of these analysis were parallelized with GNU parallel [66]. We used DeviaTE (v0.3.8)[69] to estimate the copy numbers of TEs and to visualize the abundance and the diversity of TEs. DeviaTE estimates the copy numbers of TEs in haploid genomes by normalizing the coverage of a TE sequence to the coverage of single copy genes. To estimate the number of reads mapping to each TE (reads per million mapped reads; rpm) we used PopoolationTE2 v1.10.03 [36].

To identify TE insertions in the long-read assemblies of the *D. melanogaster* strains Canton-S, Iso1, Pi2, Dgrp-732 [18, 74]) we used RepeatMasker (open-4.0.7; -no-is -s -nolow; [63]) providing the consensus sequences of TEs [52] as custom library. We merged fragmented matches using a Python script (*rm* − *defragmenter*.*py* –dist 100) and visualized the joint distribution of the insert size and the divergence using hexagonal heatmaps (ggplot2 [71]).

### UMAP and PCA

In order to identify population structure in the GDL samples we estimated the frequencies of TE-specific SNPs. The frequencies of TE-specific SNPs was inferred from reads aligned to the consensus sequences of TEs (see above). This frequency will reflect the TE composition in a given sample. For example if a specimen has 10 Opus insertions and a biallelic SNP with a frequency of 0.6 in Opus at position 351, than about 6 Opus insertions in the sample will have the SNP and 4 will not have it. We estimated the allele frequency of TE-specific SNPs in the GDL samples with DeviaTE [69]. We filtered the SNPs by solely using bi-allelic SNPs and removing SNPs solely found in few samples (≤3 samples) using a Python script (mpileup2PCA.py). These filtered SNPs were then subjected to multi-dimensional analysis in R, using UMAP (umap package; v0.2.10.0 [46]) and PCA (prcomp).

### piRNAs

We utilized data from 10 GDL strains [42] for the piRNA analysis. We removed the adaptor sequence “TG-GAATTCTCGGGTGCCAAGG” using cutadapt (v4.4 [45]) and filtered for reads having a length between 18 and 36nt. Subsequently, the reads were aligned to a database encompassing *D. melanogaster* miRNAs, mRNAs, rRNAs, snRNAs, snoRNAs, tRNAs [67], and TE sequences [52] using novoalign (v3.09.04). To compute the ping-pong signatures and visualize the piRNA abundance along the sequence of the TEs, we employed a previously developed Python scripts [58].

### Origin of the HTs

To identify potential donor species for the HT of Blood, Opus, 412, Tirant, the I-element, Hobo and the P-element we investigated the long-read assemblies of 101 diverse drosophilid species [34] and of 99 different insect species [28] (supplementary table S5). We included the long-read assemblies of a recently collected *D. melanogaster* (Pi2) and *D. simulans* (SZ232) strain into the analysis [73, 72]). The assemblies were downloaded with NCBI datasets (v14.24.0). We used RepeatMasker [63] (open-4.0.7; -no-is -s -nolow) with the consensus sequences of TEs [52] as custom library to identify TE insertions in these assemblies. A Python script was used to identify for each assembly and for each TE family the best match (i.e. the HSP with the highest alignment score) (*process* − 101*genomes*.*py*). The script further computes for each TE family the similarity of the best match to the consensus sequence as (*s* = *max*(*rms*_*i*_)*/max*(*rms*_*all*_), where *max*(*rmsi*) is the highest RepeatMasker score (rms) in a given assembly (*i*) and *max*(*rms*_*all*_) the highest score in any of the assemblies. The similarity is a value between 0 and 1, where 0 indicates no similarity to the consensus sequence of the TE and 1 a high similarity.

## Author contributions

RK conceived the work. AS, RP, FW and RK analysed the data. RK wrote the first draft. AS, RP and FW contributed to writing.

## Acknowledgments

This work was supported by an Austrian Science Fund (FWF) grant P35093 and P34965 to RK. We thank John Pool and Marcus Stensmyr for generously making the genomes of the historical specimens publicly available. We further thank John Pool for discussions. We thank all members of the Institute of Population Genetics for feedback and support.

## Data availability

All analysis performed in this work were documented in RMarkdown and have been made publicly available, together with the resulting figures, at github https://github.com/Almo96/dmel_TE_invasions (see *.md files).

## References

[1] Anxolabéhére, D., Kidwell, M. G., and Periquet, G. (1988). Molecular characteristics of diverse populations are consistent with the hypothesis of a recent invasion of Drosophila melanogaster by mobile P elements. Molecular biology and evolution, 5(3):252–69.

[2] Bartolomé, C., Bello, X., and Maside, X. (2009). Widespread evidence for horizontal transfer of transposable elements across Drosophila genomes. Genome biology, 10(2):R22.

[3] Bergman, C. M., Quesneville, H., Anxolabéhére, D., and Ashburner, M. (2006). Recurrent insertion and duplication generate networks of transposable element sequences in the Drosophila melanogaster genome. Genome biology, 7(11):R112.

[4] Blumenstiel, J. P. (2019). Birth, School, Work, Death and Resurrection: The Life Stages and Dynamics of Transposable Element Proliferation. Genes, 10(5):336.

[5] Bock, I. and Parsons, P. (1981). Species of Australia and New Zealand. In Ashburner, M., Carson, L., and Thompson, J. J., editors, The genetics and biology of Drosophila, volume 3a, pages 349–393. Academic Press, Oxford.

[6] Bonnivard, E., Bazin, C., Denis, B., and Higuet, D. (2000). A scenario for the hobo transposable element invasion, deduced from the structure of natural populations of Drosophila melanogaster using tandem TPE repeats. Genetical Research, 75(1):13–23.

[7] Bourque, G., Burns, K. H., Gehring, M., Gorbunova, V., Seluanov, A., Hammell, M., Imbeault, M., Izsvák, Z., Levin, H. L., Macfarlan, T. S., Mager, D. L., and Feschotte, C. (2018). Ten things you should know about transposable elements. pages 1–12.

[8] Bowen, N. J. and McDonald, J. F. (2001). Drosophila euchromatic LTR retrotransposons are much younger than the host species in which they reside. Genome research, 11(9):1527–1540.

[9] Brennecke, J., Aravin, A. A., Stark, A., Dus, M., Kellis, M., Sachidanandam, R., and Hannon, G. J. (2007). Discrete small RNA-generating loci as master regulators of transposon activity in Drosophila. Cell, 128(6):1089–1103.

[10] Brennecke, J., Malone, C. D., Aravin, A. A., Sachidanandam, R., Stark, A., and Hannon, G. J. (2008). An epigenetic role for maternally inherited piRNAs in transposon silencing. Science, 322(5906):1387–1392.

[11] Bucheton, A., Vaury, C., Chaboissier, M. C., Abad, P., Pélisson, A., and Simonelig, M. (1992). I elements and the Drosophila genome. Genetica, 86(1-3):175–90.

[12] Casacuberta, E. and González, J. (2013). The impact of transposable elements in environmental adaptation. Molecular ecology, 22(6):1503–17.

[13] Daniels, S. B., Chovnick, A., and Boussy, I. (1990a). Distribution of hobo transposable elements in the genus drosophila. Molecular biology and evolution, 7(6):589–606.

[14] Daniels, S. B., Peterson, K. R., Strausbaugh, L. D., Kidwell, M. G., and Chovnick, A. (1990b). Evidence for horizontal transmission of the P transposable element between Drosophila species. Genetics, 124(2):339–55.

[15] Diaz-Papkovich, A., Anderson-Trocmé, L., Ben-Eghan, C., and Gravel, S. (2019). Umap reveals cryptic population structure and phenotype heterogeneity in large genomic cohorts. PLoS genetics, 15(11):e1008432.

[16] Diaz-Papkovich, A., Anderson-Trocmé, L., and Gravel, S. (2021). A review of UMAP in population genetics. J. Hum. Genet., 66(1):85–91.

[17] Elena, S. F., Ekunwe, L., Hajela, N., Oden, S. A., and Lenski, R. E. (1998). Distribution of fitness effects caused by random insertion mutations in Escherichia coli. Genetica, 102-103:349–358.

[18] Ellison, C. E. and Cao, W. (2019). Nanopore sequencing and Hi-C scaffolding provide insight into the evolutionary dynamics of transposable elements and piRNA production in wild strains of Drosophila melanogaster. Nucleic Acids Research, pages 1–14.

[19] Engels, W. R. (1992). The origin of P elements in Drosophila melanogaster. BioEssays, 14(10):681–6.

[20] Feschotte, C. (2008). Transposable elements and the evolution of regulatory networks. Nat. Rev. Genet., 9:397–405.

[21] Finnegan, D. J. (1989). Eukaryotic transposable elements and genome evolution. Trends in Genetics, 5(4):103–107.

[22] González, J., Lenkov, K., Lipatov, M., Macpherson, J. M., and Petrov, D. A. (2008). High rate of recent transposable element–induced adaptation in Drosophila melanogaster . PLoS biology, 6(10):e251.

[23] Goriaux, C., Théron, E., Brasset, E., and Vaury, C. (2014). History of the discovery of a master locus producing piRNAs: The flamenco/COM locus in Drosophila melanogaster . Frontiers in Genetics, 5:1–8.

[24] Grenier, J. K., Arguello, J. R., Moreira, M. C., Gottipati, S., Mohammed, J., Hackett, S. R., Boughton, R., Greenberg, A. J., and Clark, A. G. (2015). Global diversity lines–a five-continent reference panel of sequenced drosophila melanogaster strains. G3: Genes, Genomes, Genetics, 5(4):593–603.

[25] Gunawardane, L. S., Saito, K., Nishida, K. M., Miyoshi, K., Kawamura, Y., Nagami, T., Siomi, H., and Siomi, M. C. (2007). A slicer-mediated mechanism for repeat-associated siRNA 5’ end formation in Drosophila. Science, 315(5818):1587–1590.

[26] Hill, T., Schlötterer, C., and Betancourt, A. J. (2016). Hybrid Dysgenesis in Drosophila simulans Associated with a Rapid Invasion of the P-Element. PLoS Genetics, 12(3):e1005920.

[27] Hoskins, R. A., Carlson, J. W., Wan, K. H., Park, S., Mendez, I., Galle, S. E., Booth, B. W., Pfeiffer, B. D., George, R. A., Svirskas, R., Krzywinski, M., Schein, J., Accardo, M. C., Damia, E., Messina, G., Méndez-Lago, M., de Pablos, B., Demakova, O. V., and E. N. Andreyeva, … S. E. Celniker (2015). The Release 6 reference sequence of the Drosophila melanogaster genome. Genome Research, 25(3):445–458.

[28] Hotaling, S., Sproul, J. S., Heckenhauer, J., Powell, A., Larracuente, A. M., Pauls, S. U., Kelley, J. L., and Frandsen, P. B. (2021). Long Reads Are Revolutionizing 20 Years of Insect Genome Sequencing. Genome Biology and Evolution, 13(8).

[29] Kapitonov, V. V. and Jurka, J. (2003). Molecular paleontology of transposable elements in the Drosophila melanogaster genome. Proceedings of the National Academy of Sciences of the United States of America, 100(11):6569–74.

[30] Kazazian, H. H. (2004). Mobile elements: drivers of genome evolution. Science, 303:1626–32.

[31] Keller, A. (2007). Drosophila melanogaster’s history as a human commensal. Current biology, 17(3):R77–R81.

[32] Kidwell, M. G. (1983). Evolution of hybrid dysgenesis determinants in Drosophila melanogaster . Proceedings of the National Academy of Sciences, 80(6):1655–1659.

[33] Kidwell, M. G., Kidwell, J. F., and Sved, J. A. (1977). Hybrid dysgenesis in Drosophila melanogaster : A syndrome of aberrant traits including mutations, sterility and male recombination. Genetics, 86(4):813–833.

[34] Kim, B. Y., Wang, J. R., Miller, D. E., Barmina, O., Delaney, E., Thompson, A., Comeault, A. A., Peede, D., D’Agostino, E. R., Pelaez, J., Aguilar, J. M., Haji, D., Matsunaga, T., Armstrong, E. E., Zych, M., Ogawa, Y., Stamenković-Radak, M., Jelić, M., Veselinović, M. S., Tanasković, M., Erić, P., Gao, J.-J., Katoh, T. K., Toda, M. J., Watabe, H., Watada, M., Davis, J. S., Moyle, L. C., Manoli, G., Bertolini, E., Koštál, V., Hawley, R. S., Takahashi, A., Jones, C. D., Price, D. K., Whiteman, N., Kopp, A., Matute, D. R., and Petrov, D. A. (2021). Highly contiguous assemblies of 101 drosophilid genomes. eLife, 10:e66405.

[35] Kofler, R., Betancourt, A. J., and Schlötterer, C. (2012). Sequencing of Pooled DNA Samples (Pool-Seq) Uncovers Complex Dynamics of Transposable Element Insertions in Drosophila melanogaster. PLoS genetics, 8(1):e1002487.

[36] Kofler, R., Gomez-Sanchez, D., and Schlötterer, C. (2016). PoPoolationTE2: Comparative Population Genomics of Transposable Elements Using Pool-Seq. MBE, 33(10):2759–2764.

[37] Kofler, R., Hill, T., Nolte, V., Betancourt, A., and Schlötterer, C. (2015a). The recent invasion of natural Drosophila simulans populations by the P-element. PNAS, 112(21):6659–6663.

[38] Kofler, R., Nolte, V., and Schlötterer, C. (2015b). Tempo and mode of transposable element activity in Drosophila. PLoS Genetics, 11(7):e1005406.

[39] Le Thomas, A., Rogers, A. K., Webster, A., Marinov, G. K., Liao, S. E., Perkins, E. M., Hur, J. K., Aravin, A. A., and Tóth, K. F. (2013). Piwi induces piRNA-guided transcriptional silencing and establishment of a repressive chromatin state. Genes and Development, 27(4):390–399.

[40] Li, H. and Durbin, R. (2009). Fast and accurate short read alignment with Burrows–Wheeler transform. Bioinformatics, 25(14):1754–1760.

[41] Loreto, E. L. S., Carareto, C. M. A., and Capy, P. (2008). Revisiting horizontal transfer of transposable elements in Drosophila. Heredity, 100(6):545–54.

[42] Luo, S., Zhang, H., Duan, Y., Yao, X., Clark, A. G., and Lu, J. (2020). The evolutionary arms race between transposable elements and pirnas in drosophila melanogaster. BMC Evolutionary Biology, 20(1):1–18.

[43] Malone, C. D., Brennecke, J., Dus, M., Stark, A., McCombie, W. R., Sachidanandam, R., and Hannon, G. J. (2009). Specialized piRNA pathways act in germline and somatic tissues of the Drosophila ovary. Cell, 137(3):522–535.

[44] Marí-Ordóñez, A., Marchais, A., Etcheverry, M., Martin, A., Colot, V., and Voinnet, O. (2013). Reconstructing de novo silencing of an active plant retrotransposon. Nature genetics, 45(9):1029–39.

[45] Martin, M. (2011). Cutadapt removes adapter sequences from high-throughput sequencing reads. EMBnet. journal, 17(1):pp–10.

[46] McInnes, L., Healy, J., and Melville, J. (2018). Umap: Uniform manifold approximation and projection for dimension reduction.

[47] Moon, S., Cassani, M., Lin, Y. A., Wang, L., Dou, K., and Zhang, Z. Z. (2018). A Robust Transposon-Endogenizing Response from Germline Stem Cells. Developmental Cell, 47(5):660–671.e3.

[48] Ozata, D. M., Gainetdinov, I., Zoch, A., O’Carroll, D., and Zamore, P. D. (2018). PIWI-interacting RNAs: small RNAs with big functions. Nature Reviews Genetics, 20(2):89–108.

[49] Pasyukova, E., Nuzhdin, S., Morozova, T., and Mackay, T. (2004). Accumulation of TEs in the genome of D. melanogaster is associated with a decrease in fitness. Journal of Heredity, 95(4):284–290.

[50] Peccoud, J., Loiseau, V., Cordaux, and Gilbert, C. (2017). Massive horizontal transfer of transposable elements in insects. PNAS, 114(18):4721–26.

[51] Periquet, G., Hamelin, M. H., Bigot, Y., and Lepissier, A. (1989). Geographical and historical patterns of distribution of hobo elements in drosophila melanogaster populations. Journal of Evolutionary Biology, 2(3):223–229.

[52] Quesneville, H., Bergman, C. M., Andrieu, O., Autard, D., Nouaud, D., Ashburner, M., and Anxolabehere, D. (2005). Combined evidence annotation of transposable elements in genome sequences. PLoS Comp. Biol., 1(2):166–175.

[53] Robertson, H. M. (1993). The mariner transposable element is widespread in insects. Nature, 362(6417):241–5.

[54] Said, I., McGurk, M. P., Clark, A. G., and Barbash, D. A. (2021). Patterns of piRNA Regulation in Drosophila Revealed through Transposable Element Clade Inference. Molecular Biology and Evolution, 39(1).

[55] Sánchez-Gracia, A., Maside, X., and Charlesworth, B. (2005). High rate of horizontal transfer of transposable elements in drosophila. Trends in genetics, 21(4):200–203.

[56] Sarkies, P., Selkirk, M. E., Jones, J. T., Blok, V., Boothby, T., Goldstein, B., Hanelt, B., Ardila-Garcia, A., Fast, N. M., Schiffer, P. M., Kraus, C., Taylor, M. J., Koutsovoulos, G., Blaxter, M. L., and Miska, E. A. (2015). Ancient and novel small RNA pathways compensate for the loss of piRNAs in multiple independent nematode lineages. PLoS Biol., 13(2):1–20.

[57] Schwarz, F., Wierzbicki, F., Senti, K.-A., and Kofler, R. (2020). Tirant stealthily invaded natural drosophila melanogaster populations during the last century. bioRxiv.

[58] Selvaraju, D., Wierzbicki, F., and Kofler, R. (2022). P-element invasions in Drosophila erecta shed light on the establishment of host control over a transposable element. bioRxiv.

[59] Shpak, M., Ghanavi, H. R., Lange, J. D., Pool, J. E., and Stensmyr, M. C. (2023). Genomes from 25 historical Drosophila melanogaster specimens illuminate adaptive and demographic changes across more than 200 years of evolution. bioRxiv.

[60] Sienski, G., Dönertas, D., and Brennecke, J. (2012). Transcriptional silencing of transposons by Piwi and maelstrom and its impact on chromatin state and gene expression. Cell, 151(5):964–980.

[61] Simmons, G. (1992). Horizontal transfer of hobo transposable elements within the drosophila melanogaster species complex: evidence from dna sequencing. Molecular biology and evolution, 9(6):1050–1060.

[62] Singh, N. D. and Petrov, D. A. (2004). Rapid sequence turnover at an intergenic locus in Drosophila. Molecular biology and evolution, 21(4):670–80.

[63] Smit, A. F. A., Hubley, R., and Green, P. (1996). RepeatMasker Open-3.0.

[64] Sprengelmeyer, Q. D., Mansourian, S., Lange, J. D., Matute, D. R., Cooper, B. S., Jirle, E. V., Stensmyr, M. C., and Pool, J. E. (2020). Recurrent collection of drosophila melanogaster from wild african environments and genomic insights into species history. Molecular biology and evolution, 37(3):627–638.

[65] Srivastav, S., Feschotte, C., and Clark, A. G. (2023). Rapid evolution of pirna clusters in the drosophila melanogaster ovary. bioRxiv.

[66] Tange, O. (2018). GNU parallel 2018. Lulu. com.

[67] Thurmond, J., Goodman, J. L., Strelets, V. B., Attrill, H., Gramates, L. S., Marygold, S. J., Matthews, B. B., Millburn, G., Antonazzo, G., Trovisco, V., et al. (2019). Flybase 2.0: the next generation. Nucleic acids research, 47(D1):D759–D765.

[68] Vieira, C., Lepetit, D., Dumont, S., and Biémont, C. (1999). Wake up of transposable elements following Drosophila simulans worldwide colonization. Molecular biology and evolution, 16(9):1251–5.

[69] Weilguny, L. and Kofler, R. (2019). DeviaTE: Assembly-free analysis and visualization of mobile genetic element composition. Molecular ecology resources, 19(5):1346–1354.

[70] Wicker, T., Sabot, F., Hua-Van, A., Bennetzen, J. L., Capy, P., Chalhoub, B., Flavell, A., Leroy, P., Morgante, M., Panaud, O., et al. (2007). A unified classification system for eukaryotic transposable elements. Nature Reviews Genetics, 8(12):973–982.

[71] Wickham, H. (2016). ggplot2: Elegant Graphics for Data Analysis. Springer-Verlag New York.

[72] Wierzbicki, F. and Kofler, R. (2023). The composition of piRNA clusters in Drosophila melanogaster deviates from expectations under the trap model. bioRxiv.

[73] Wierzbicki, F., Kofler, R., and Signor, S. (2023). Evolutionary dynamics of piRNA clusters in Drosophila. Molecular Ecology, 32:1306–1322.

[74] Wierzbicki, F., Schwarz, F., Cannalonga, O., and Kofler, R. (2021). Novel quality metrics allow identifying and generating high-quality assemblies of pirna clusters. Molecular Ecology Resources.

[75] Yamanaka, S., Siomi, M. C., and Siomi, H. (2014). piRNA clusters and open chromatin structure. Mobile DNA, 5(1):22.

[76] Zanni, V., Eymery, A., Coiffet, M., Zytnicki, M., Luyten, I., Quesneville, H., Vaury, C., and Jensen, S. (2013). Distribution, evolution, and diversity of retrotransposons at the flamenco locus reflect the regulatory properties of piRNA clusters. PNAS, 110(49):19842–19847.

