## Supplement for "Genomes of historical specimens reveal multiple invasions of LTR retrotransposons in *Drosophila melanogaster* populations during the 19^*th*^ century"

#### Supplementary figures

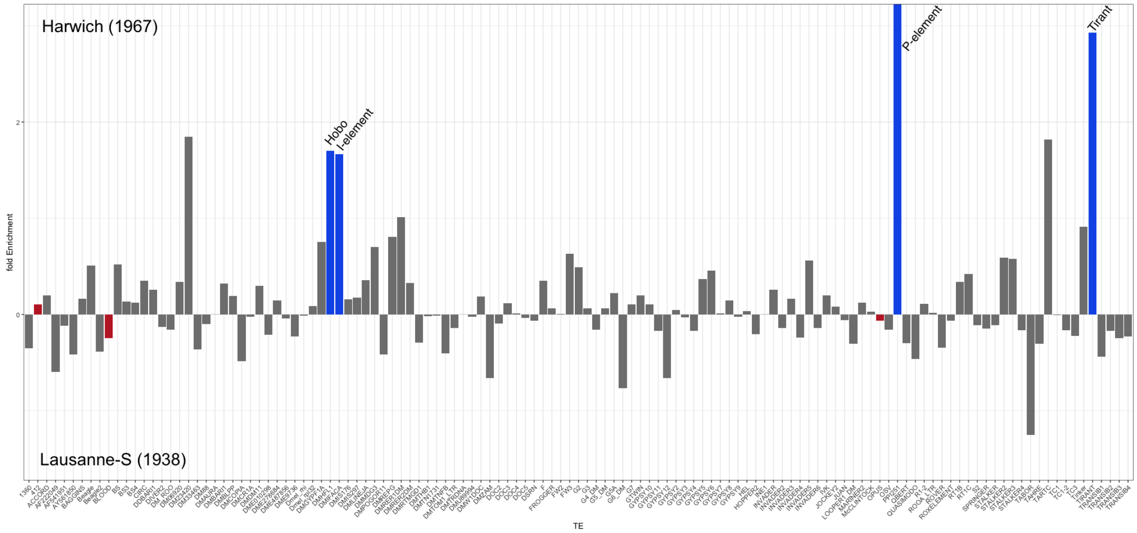


Figure 1: Differences in TE abundance between a strain collected in 1967 (Harwich) and 1938 (Lausanne-S). For each TE family (x-axis) we show the difference in TE copy numbers per haploid genome as estimated with DeviaTE (y-axis). As expected Hobo, the I-element, the P-element and Tirant (blue bars) are overrepresented in Harwich [Schwarz et al., 2020].

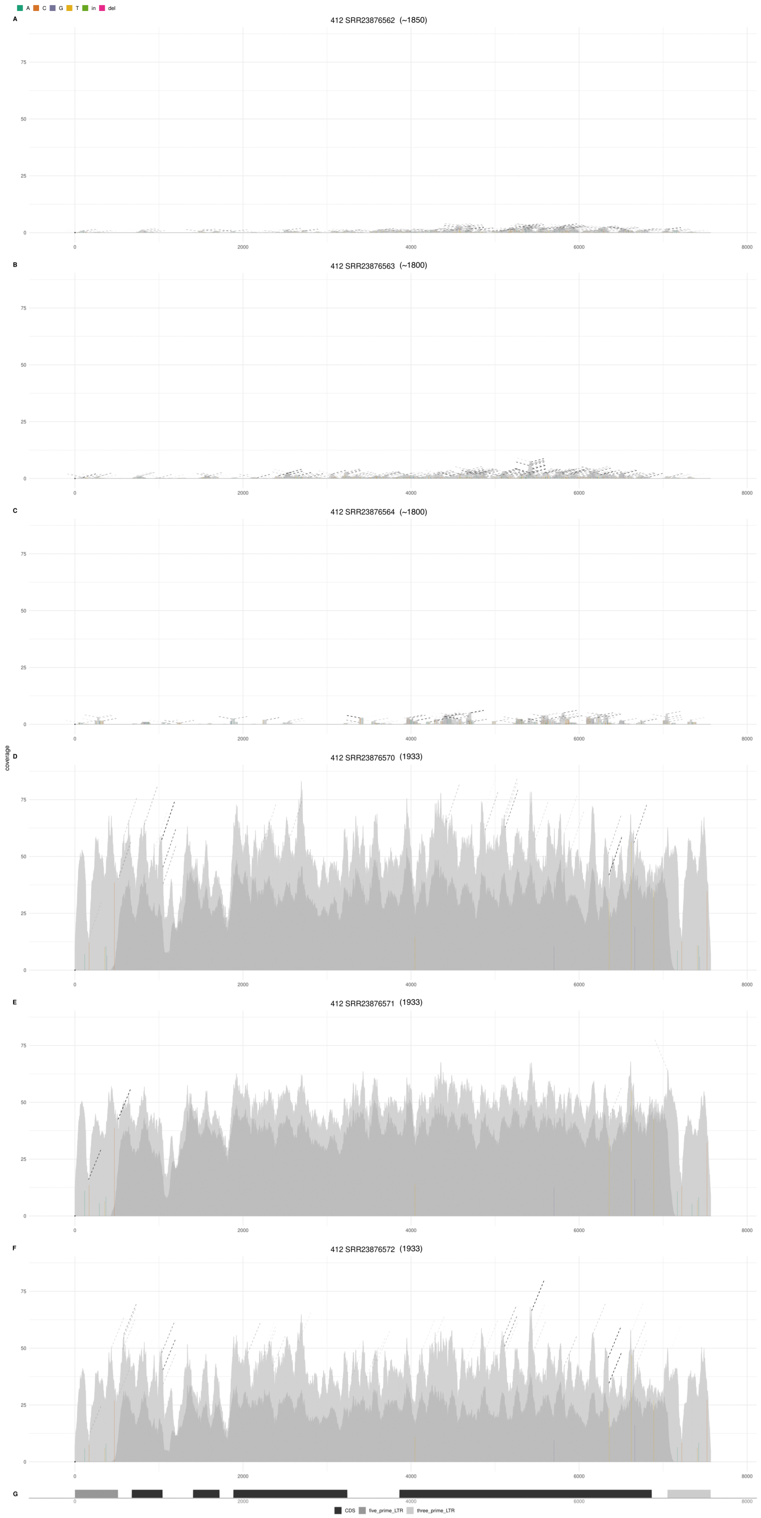


Figure 2: DeviaTE plots for 412 using three strains sampled around 1800 (top three) and three strains sampled around 1933 (bottom three). The coverage of 412 was normalized to the coverage of single-copy genes. Single-nucleotide polymorphisms (SNPs) and small internal deletions (indels) are shown as colored lines. The coverage based on uniquely and ambiguously mapped reads is shown in dark and light gray, respectively.

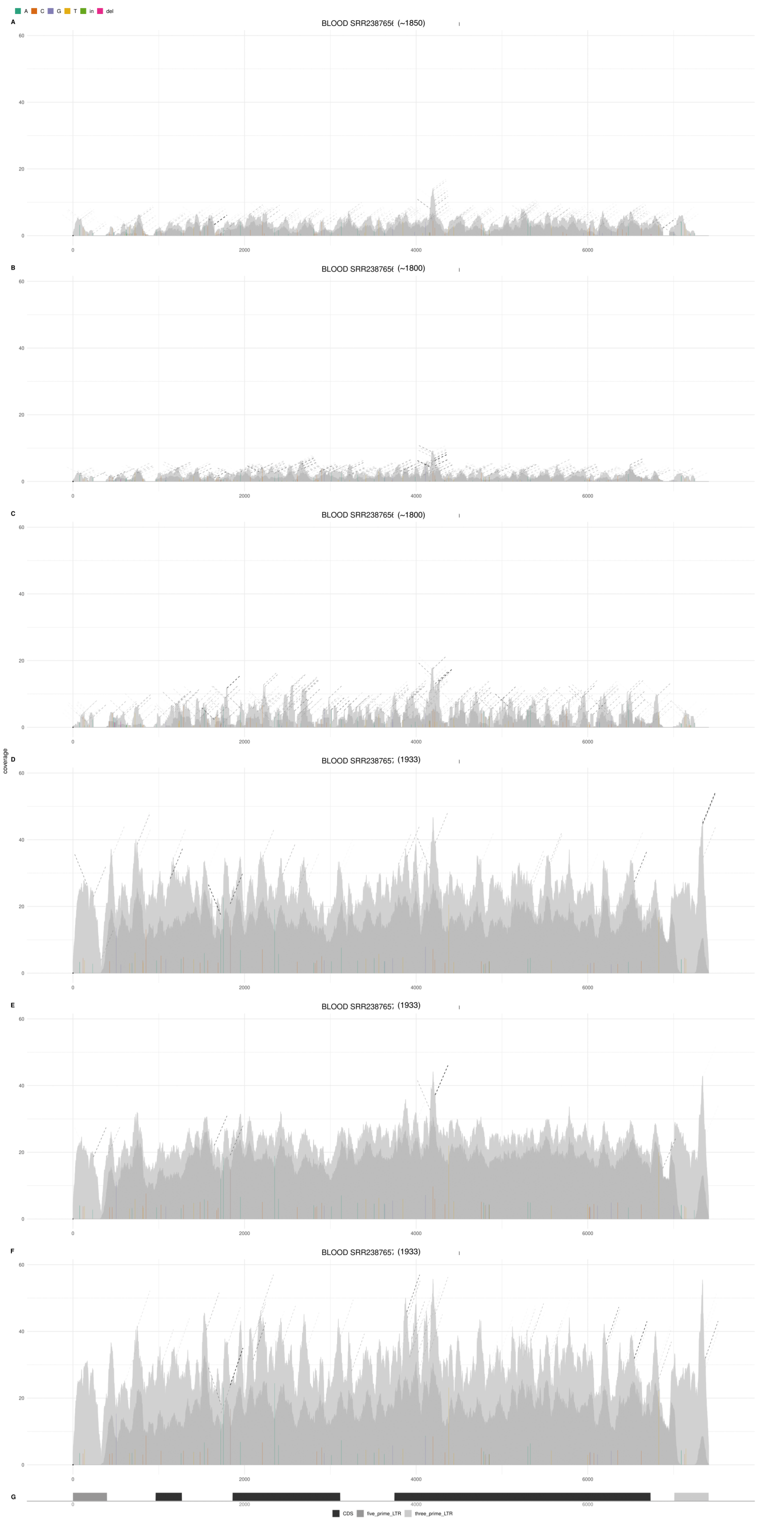


Figure 3: DeviaTE plots for Blood using three strains sampled around 1800 (top three) and three strains sampled around 1933 (bottom three). The coverage of Blood was normalized to the coverage of single-copy genes. Single-nucleotide polymorphisms (SNPs) and small internal deletions (indels) are shown as colored lines. The coverage based on uniquely and ambiguously mapped reads is shown in dark and light gray, respectively.

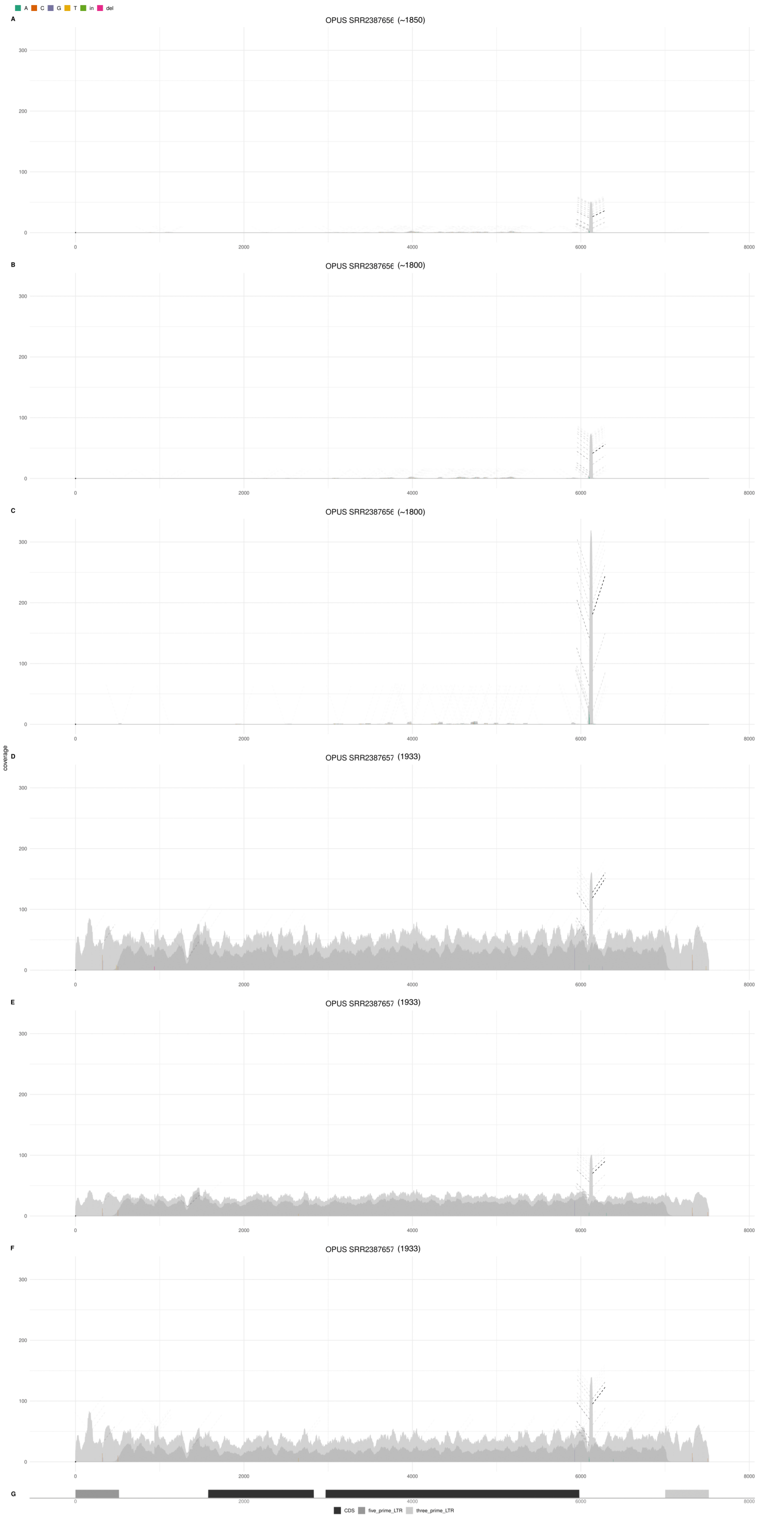


Figure 4: DeviaTE plots for Opus using three strains sampled around 1800 (top three) and three strains sampled around 1933 (bottom three). The coverage of Opus was normalized to the coverage of single-copy genes. Single-nucleotide polymorphisms (SNPs) and small internal deletions (indels) are shown as colored lines. The coverage based on uniquely and ambiguously mapped reads is shown in dark and light gray, respectively.

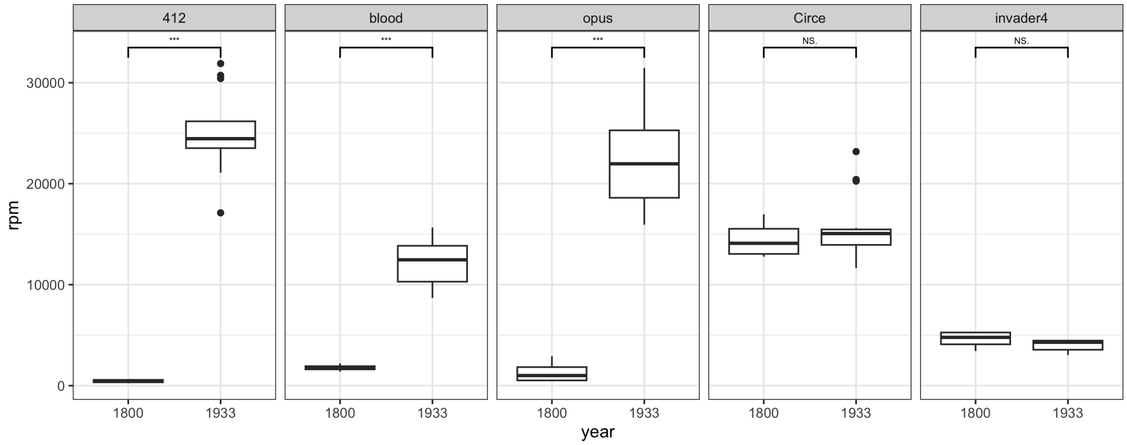


Figure 5: Abundance of 412, Opus and Blood in historical specimens collected around 1800 (6 samples) and 1933 (16 samples). As controls Circe and Invader-4 are included. The abundance of TEs is provided in reads mapping to the TE out of a million mapped reads (rpm) and the significance was computed with Wilcoxon rank sum tests.

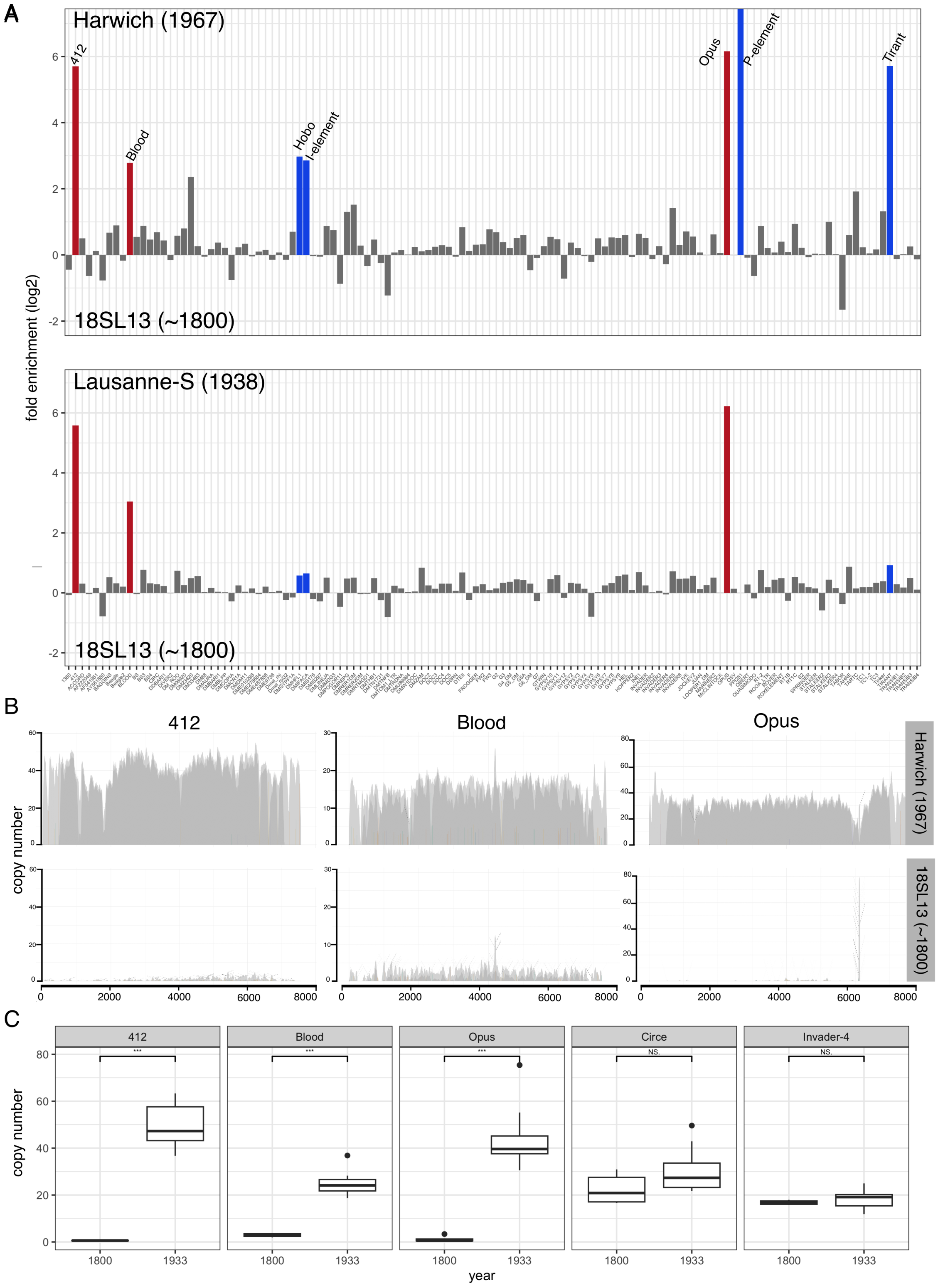


Figure 6: Reproducing figure 1 of the main manuscript with reads of length 50 (reads with a length of 100bp were used for the figure in the main manuscript). Note that the analysis of reads with 50bp length confirms that Blood, 412 and Opus have highly elevated copy numbers in strains sampled $\geq$ 1933 as compared to strains sampled around 1800.

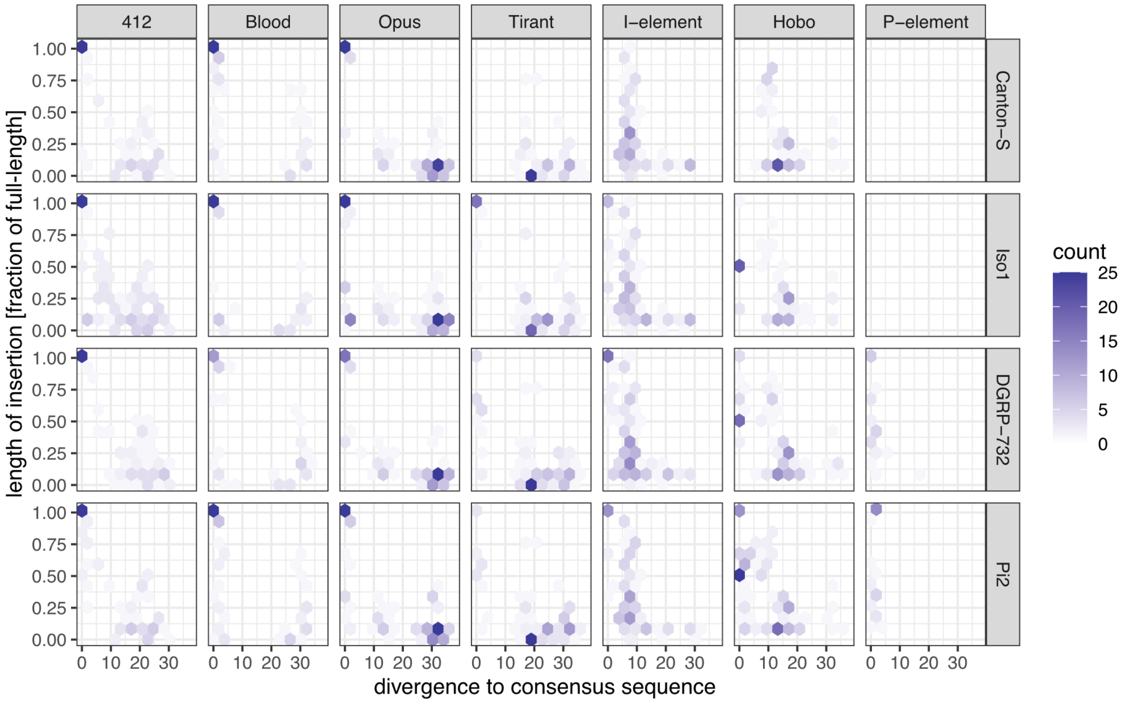


Figure 7: 2D-histograms showing the distribution of TE insertions in long-read assemblies of four different *D. melanogaster* strains. We plotted the length of the TE insertions (normalized to the full-length element) versus the divergence of the insertions to the consensus sequence. Recent insertions of a TE family are likely full-length and show little divergence (upper left corner), whereas older insertions are likely degraded and fragmented (lower right corner). Blood, Opus, and 412 may thus have recent insertions in all four analysed *D. melanogaster* strains. Note that Canton-S is an old lab strain and does therefore not have recent insertions of Tirant, I-element, Hobo and the P-element. As expected, Iso1 is lacking P-element insertions.

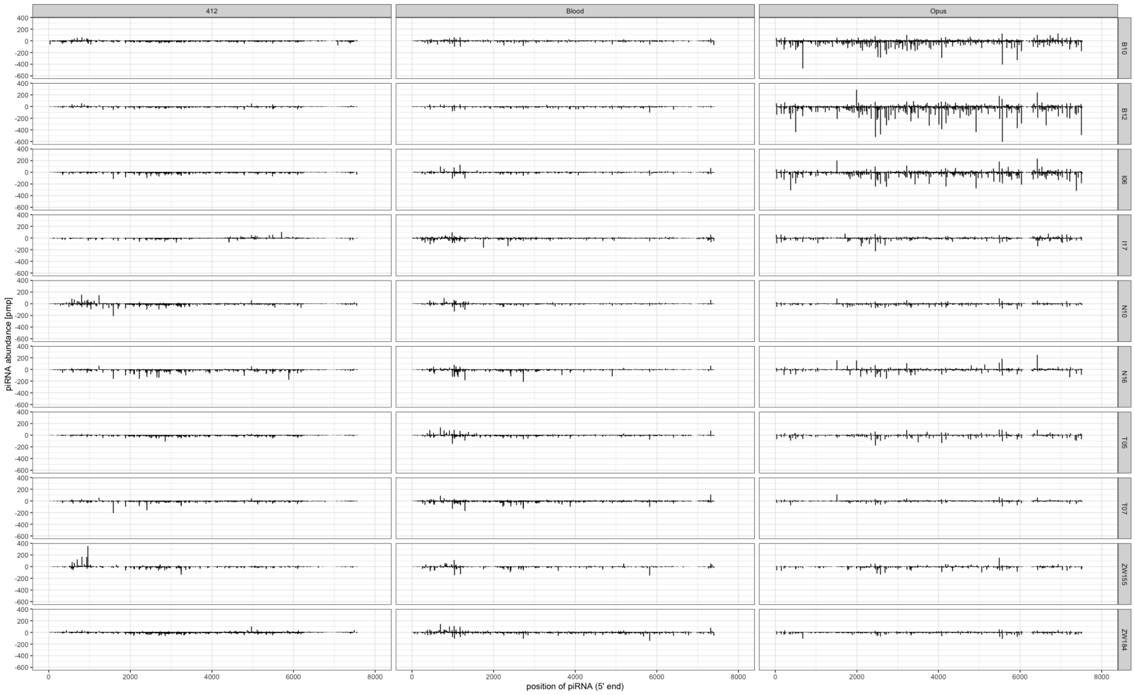


Figure 8: Distribution of piRNAs along Blood, Opus and 412 in 10 lines sampled from diverse geographic regions (GDL [Luo et al., 2020]). For each piRNA solely the 5' position is shown. Sense piRNAs are shown on the positive y-axis and antisense piRNAs on the negative y-axis. The total abundance of piRNAs was normalized to one million piRNAs.

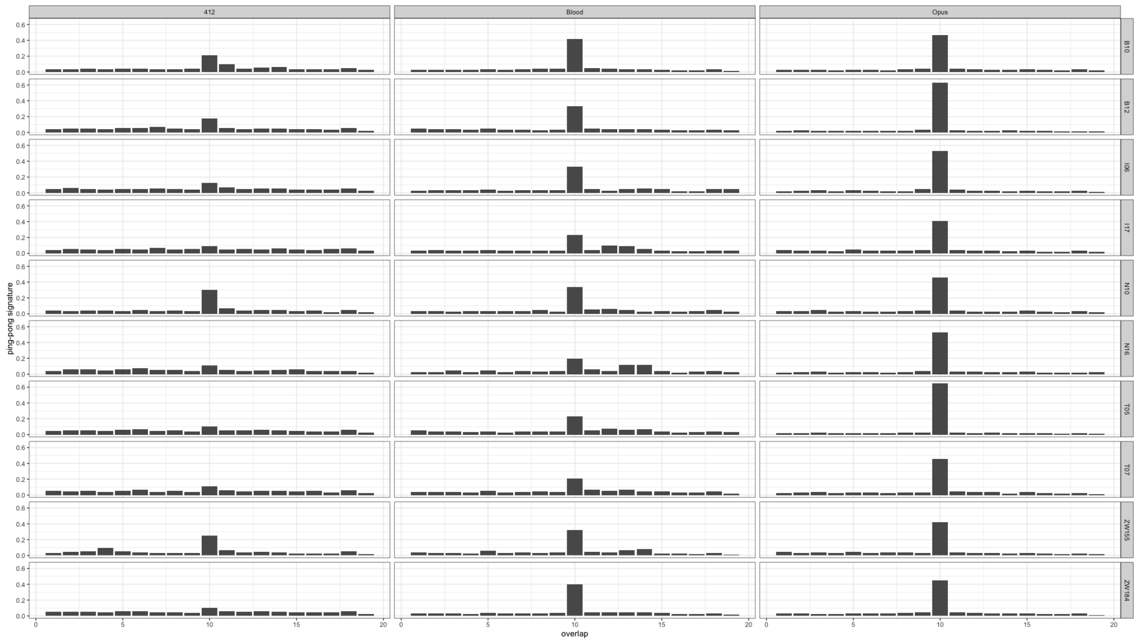


Figure 9: Ping-pong signatures for Blood, Opus and 412 in 10 lines sampled from diverse geographic regions (GDL [Luo et al., 2020]). These histograms show the distance between the 5' position of sense and antisense piRNAs. A peak at position 10 suggests that the ping-pong cycle is active and thus that the TE is silenced by the host defence.

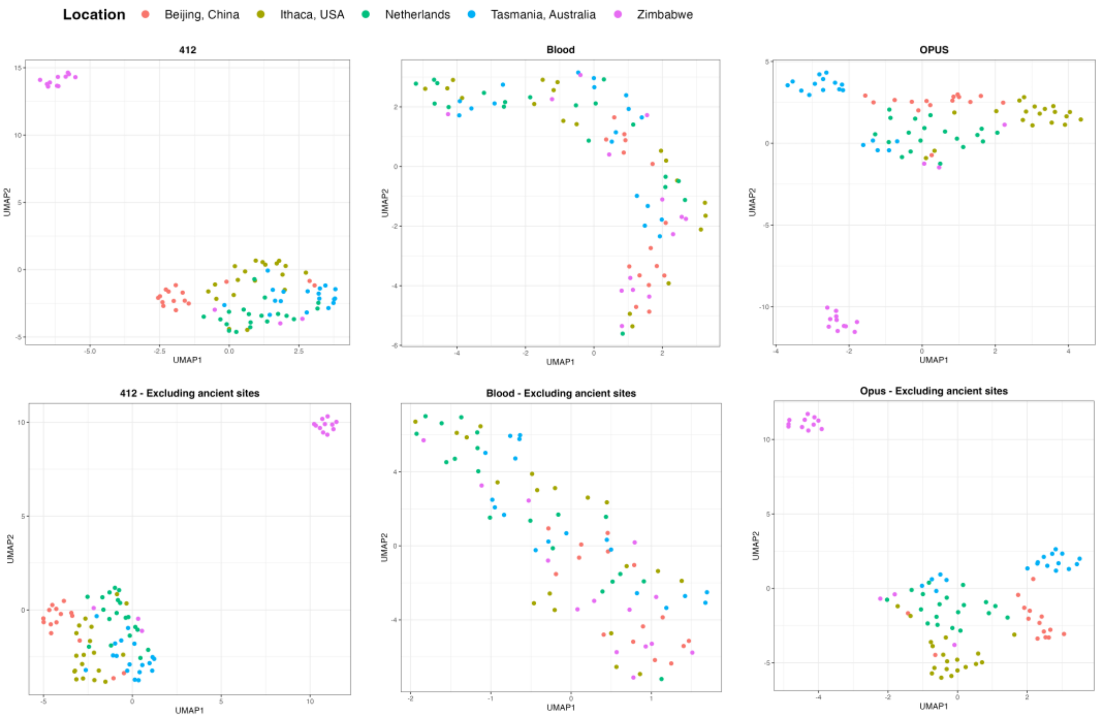


Figure 10: UMAPs for Blood, Opus and 412 in the 85 GDL strains including and excluding sites covered by ancient/degraded fragments of these TEs. We treated sites having a coverage in specimens collected around 1800 as ancient sites. Note that excluding the ancient sites only had a minor impact on the overall pattern seen in the UMAPs.

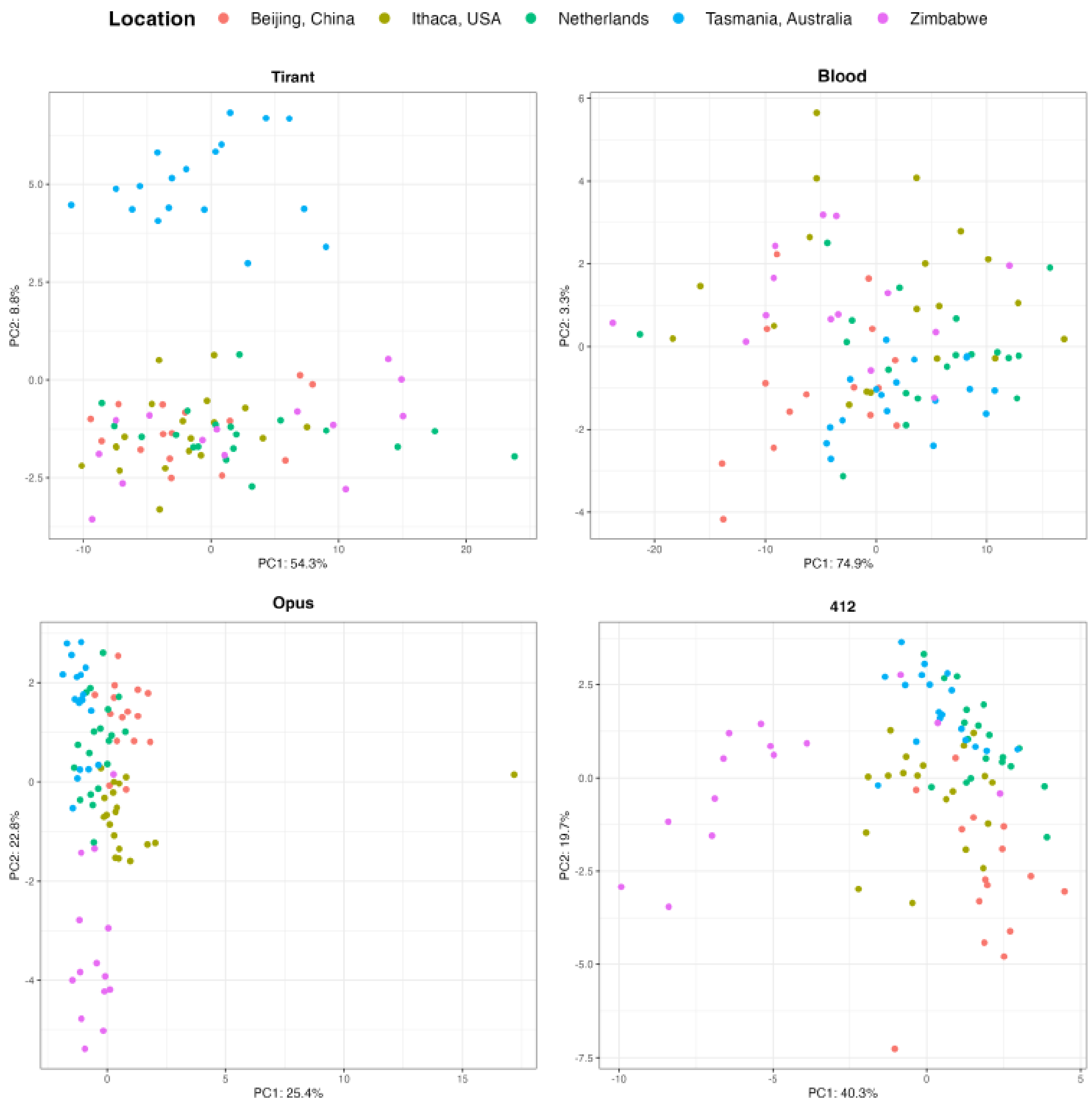


Figure 11: PCA performed with SNPs found in the TE sequences among the 85 GDL specimens.

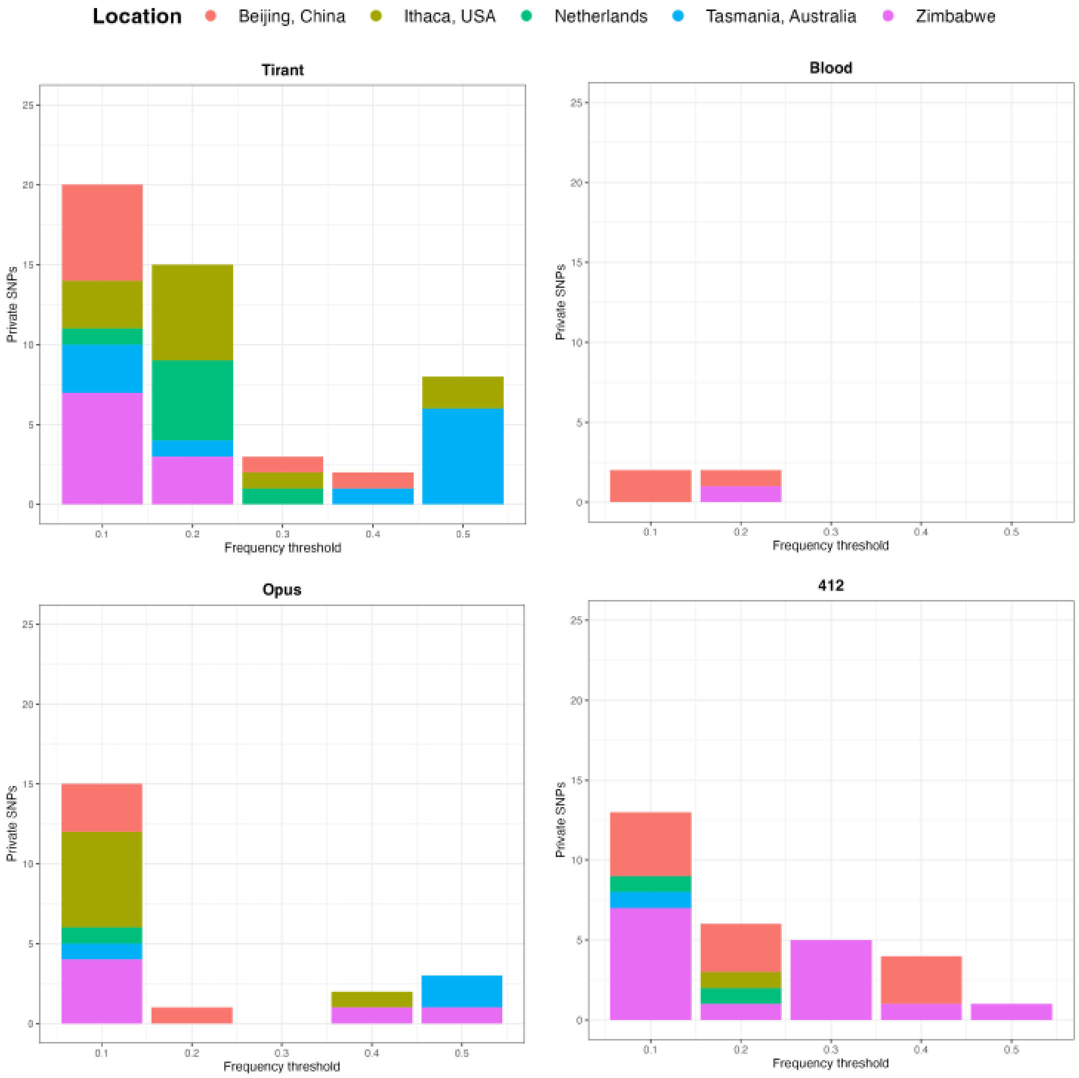


Figure 12: Frequency of diagnostic SNPs in the 85 GDL strains. We defined diagnostic SNPs to be abundant ($\geq$50%) in a population of interest but rare in all other populations (<21%). The x-axis refers to the frequency of the diagnostic SNP in a GDL strain. For example with a frequency of 0.4 about 40% of the TEs in a strain carry the diagnostic SNP. Data are shown for Tirant, Blood, Opus and 412. Notably the abundance of the diagnostic SNP reflects the clusters observed in the UMAP (main manuscript).

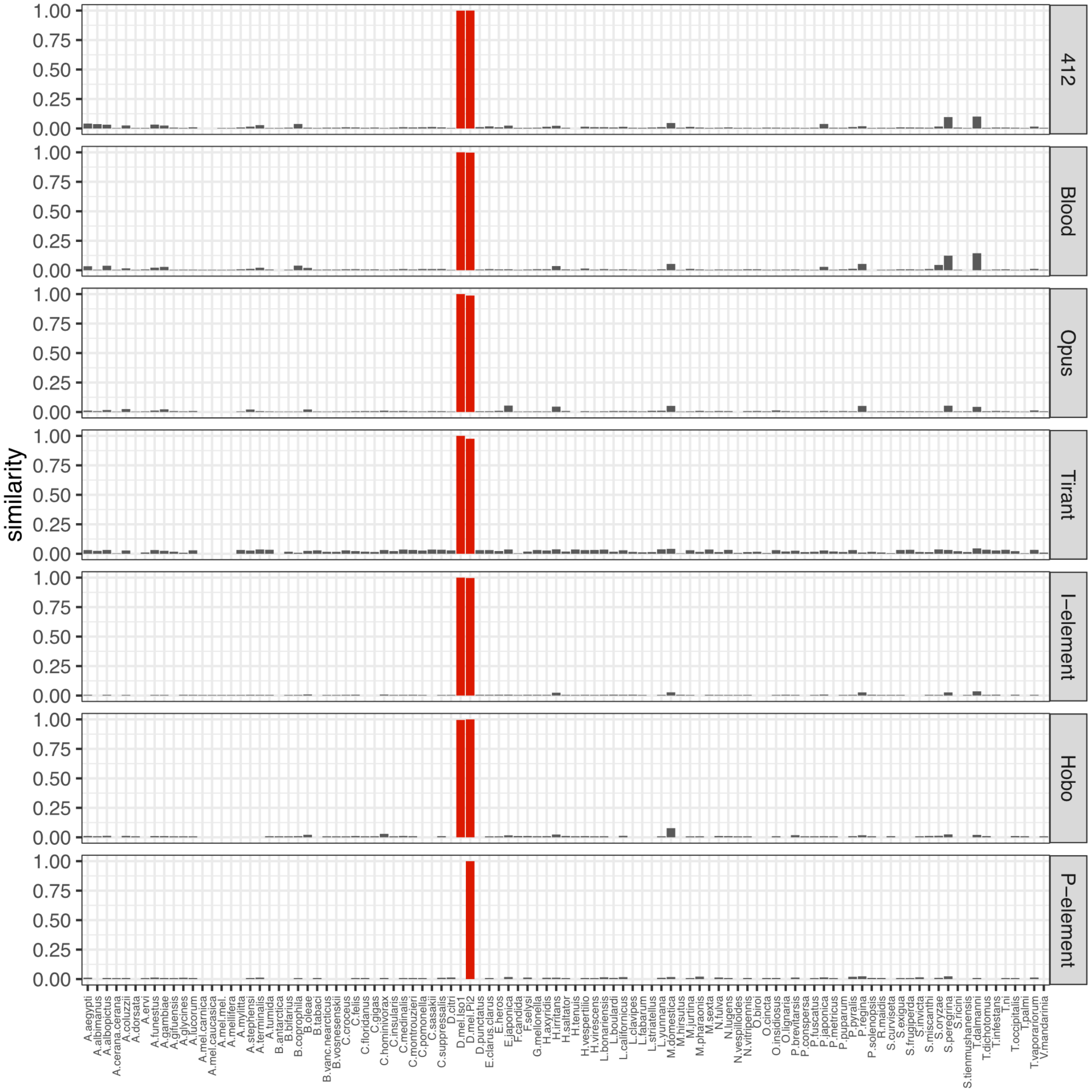


Figure 13: Barplots show the similarity of TE insertions in a given assembly to the consensus sequence of a TE for 99 long-read assemblies of diverse insect species. As reference, *D. melanogaster* (red) is included. For example, 0.9 means that at least one TE insertion in a given assembly has a high similarity ($\approx 90\%$) to the consensus sequence of the TE.

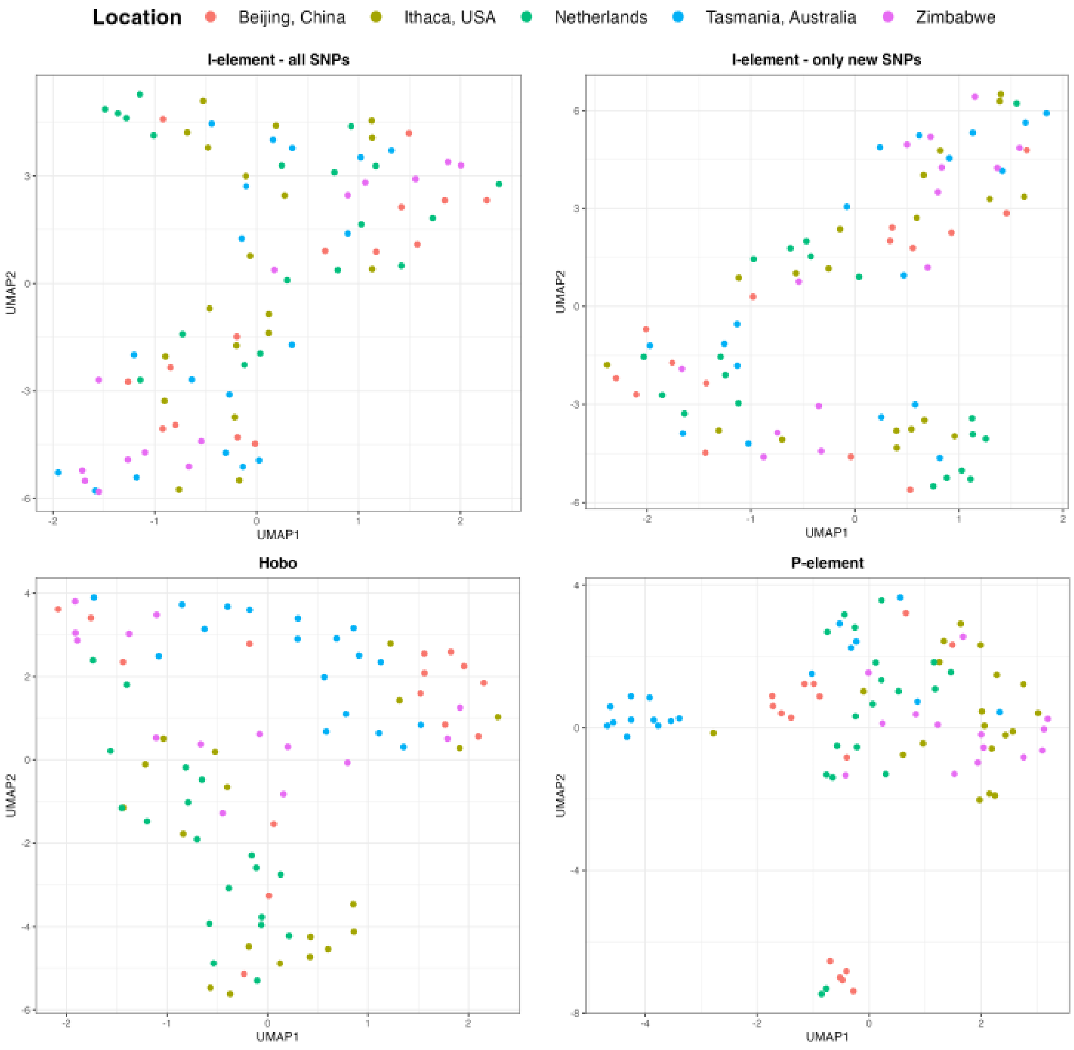


Figure 14: UMAPs for the I-element, Hobo and the P-element in the 85 GDL strains. UMAPs are based on the allele frequency of TE specific SNPs. The ancient I-element led to many diverse insertions which accumulated a substantial amount of SNPs. As these ancient SNPs could confound an analysis of the populations structure we performed UMAPs with and without the ancient SNPs for the I-element.

#### Supplementary tables

Table 1: Overview of the TE composition in the strains used for reconstructing the history of TE invasions in *D. melanogaster* during the last 200 years. For each strain we show the SRA number, the approximate year and location of the collection and the abundance of the seven TEs that recently invaded *D. melanogaster* populations. The numbers refer to TE copy numbers per haploid genome (estimated with DeviaTE [Weilguny and Kofler, 2019]. We classified the TE content into three distinct categories: red absence of any TE sequence, yellow solely degraded TE sequences are present, green non-degraded TE sequences with a high similarity to the consensus sequence are present.

| Sample | Year | Location | 412 | Blood | Opus | Tirant | Hobo | I-ele | P-ele |
|---|---|---|---|---|---|---|---|---|---|
| SRR23876563 | 1800 | Lund, Sweden | 0.82 | 2.03 | 0.52 | 0.12 | 1.17 | 8.4 | 0 |
| SRR23876564 | 1800 | Lund, Sweden | 0.68 | 3.93 | 1.64 | 0.14 | 2.15 | 9.22 | 0 |
| SRR23876566 | 1800 | Småland, Sweden | 0.57 | 3.96 | 3.94 | 0.1 | 1.78 | 9.66 | 0 |
| SRR23876582 | 1800 | Lund, Sweden | 0.95 | 2.43 | 0.49 | 0.16 | 1.42 | 8.01 | 0 |
| SRR23876583 | 1800 | Lund, Sweden | 0.55 | 3.11 | 0.77 | 0.11 | 1.87 | 7.97 | 0 |
| SRR23876584 | 1800 | Lund, Sweden | 0.85 | 2.82 | 0.46 | 0.12 | 1.23 | 7.65 | 0 |
| SRR23876562 | 1850 | Passau, Germany | 0.55 | 3.41 | 0.38 | 0.15 | 1.88 | 7.97 | 0 |
| SRR23876569 | 1850 | Passau, Germany | 0.66 | 5.5 | 0.44 | 0.15 | 2.54 | 9.43 | 0 |
| SRR23876565 | 1875 | Zealand, Denmark | 17.12 | 14.67 | 18.92 | 4.12 | 1.74 | 9.84 | 0 |
| SRR23876567 | 1933 | Lund, Sweden | 41.65 | 23.89 | 37.73 | 0.39 | 1.59 | 8.01 | 0 |
| SRR23876568 | 1933 | Lund, Sweden | 52.08 | 28.06 | 39.39 | 0.19 | 1.31 | 10 | 0 |
| SRR23876570 | 1933 | Lund, Sweden | 48.62 | 24.01 | 51.88 | 0.48 | 2.51 | 11.58 | 0 |
| SRR23876571 | 1933 | Lund, Sweden | 46.2 | 22.9 | 30.38 | 0.29 | 2.32 | 13.77 | 0 |
| SRR23876572 | 1933 | Lund, Sweden | 37.5 | 28.72 | 37.29 | 9.61 | 2.6 | 11.4 | 0 |
| SRR23876573 | 1933 | Lund, Sweden | 64.32 | 37.22 | 74.88 | 0.44 | 1.71 | 12.36 | 0 |
| SRR23876574 | 1933 | Lund, Sweden | 49.59 | 21.03 | 43.02 | 0.16 | 1.21 | 10.94 | 0 |
| SRR23876575 | 1933 | Lund, Sweden | 48.36 | 25.1 | 33.6 | 0.18 | 2.26 | 11.29 | 0 |
| SRR23876576 | 1933 | Lund, Sweden | 45.24 | 19.19 | 33.9 | 0.2 | 1.32 | 12 | 0 |
| SRR23876577 | 1933 | Lund, Sweden | 43.07 | 26.75 | 42.7 | 0.27 | 2.19 | 11.23 | 0 |
| SRR23876578 | 1933 | Lund, Sweden | 61.01 | 27.65 | 51.32 | 0.21 | 1.52 | 11.54 | 0 |
| SRR23876579 | 1933 | Lund, Sweden | 59.49 | 22.78 | 39.25 | 0.2 | 1.27 | 13.02 | 0 |
| SRR23876580 | 1933 | Lund, Sweden | 52.27 | 22.16 | 55.14 | 0.19 | 1.51 | 10.68 | 0 |
| SRR23876581 | 1933 | Lund, Sweden | 61.07 | 21.65 | 40.02 | 0.16 | 1.41 | 10.52 | 0 |
| SRR23876585 | 1933 | Lund, Sweden | 43.24 | 25.43 | 41.77 | 0.18 | 2.15 | 9.31 | 0 |
| SRR23876586 | 1933 | Lund, Sweden | 61.87 | 27.06 | 37.55 | 0.17 | 1.37 | 11.25 | 0 |
| SRR11846555 | 1936 | Crimea, Ukraine | 46.74 | 32.89 | 30.62 | 0.65 | 4.02 | 16.45 | 0 |
| SRR11846562 | 1938 | Illinois, USA | 39.23 | 24.55 | 39.59 | 2.82 | 4.94 | 19.6 | 0 |
| SRR11846563 | 1938 | Stockholm, Sweden | 41.33 | 25.64 | 39.4 | 0.69 | 23.2 | 35.32 | 0.27 |
| SRR11846564 | 1938 | Wisconsin, USA | 39.09 | 19.05 | 35.01 | 0.7 | 3.29 | 16.89 | 0 |
| SRR11846556 | 1950 | Berlin, Germany | 34.9 | 17.91 | 26.61 | 7.46 | 3.16 | 30 | 0 |
| SRR11846554 | 1952 | Florida, USA | 38.18 | 18.12 | 34.26 | 8.75 | 20.21 | 29.16 | 42.71 |
| SRR11846561 | 1954 | Israel | 46.97 | 33.34 | 25.89 | 12.29 | 15.57 | 38.11 | 0 |
| SRR11846565 | 1959 | Japan | 41.15 | 12.45 | 28.95 | 6.84 | 5.14 | 16.05 | 0 |
| SRR11846560 | 1967 | Massachussets, USA | 42.01 | 16.06 | 33.49 | 5.33 | 10.68 | 53.68 | 47.55 |
| SRR11846559 | 1987 | - | 58.37 | 21.73 | 31.75 | 3.83 | 35.68 | 38.70 | 0 |

Table 2: Overview of the GDL strains analysed in this work. For each strain we show the accession number and the location of collection.

| **Run accession** | **Location** | **Run accession** | **Location** |
|---|---|---|---|
| SRR1662283 | Beijing, China | SRR1663570 | Netherlands |
| SRR1663528 | Beijing, China | SRR1663571 | Netherlands |
| SRR1663529 | Beijing, China | SRR1663572 | Netherlands |
| SRR1663530 | Beijing, China | SRR1663573 | Netherlands |
| SRR1663531 | Beijing, China | SRR1663574 | Netherlands |
| SRR1663532 | Beijing, China | SRR1663575 | Netherlands |
| SRR1663533 | Beijing, China | SRR1663576 | Netherlands |
| SRR1663534 | Beijing, China | SRR1663577 | Netherlands |
| SRR1663535 | Beijing, China | SRR1663578 | Netherlands |
| SRR1663536 | Beijing, China | SRR1663579 | Netherlands |
| SRR1663537 | Beijing, China | SRR1663580 | Tasmania, Australia |
| SRR1663538 | Beijing, China | SRR1663581 | Tasmania, Australia |
| SRR1663539 | Beijing, China | SRR1663582 | Tasmania, Australia |
| SRR1663540 | Beijing, China | SRR1663583 | Tasmania, Australia |
| SRR1663541 | Beijing, China | SRR1663584 | Tasmania, Australia |
| SRR1663542 | Ithaca, USA | SRR1663585 | Tasmania, Australia |
| SRR1663543 | Ithaca, USA | SRR1663586 | Tasmania, Australia |
| SRR1663544 | Ithaca, USA | SRR1663587 | Tasmania, Australia |
| SRR1663545 | Ithaca, USA | SRR1663588 | Tasmania, Australia |
| SRR1663546 | Ithaca, USA | SRR1663589 | Tasmania, Australia |
| SRR1663547 | Ithaca, USA | SRR1663590 | Tasmania, Australia |
| SRR1663548 | Ithaca, USA | SRR1663591 | Tasmania, Australia |
| SRR1663549 | Ithaca, USA | SRR1663592 | Tasmania, Australia |
| SRR1663550 | Ithaca, USA | SRR1663593 | Tasmania, Australia |
| SRR1663551 | Ithaca, USA | SRR1663594 | Tasmania, Australia |
| SRR1663552 | Ithaca, USA | SRR1663595 | Tasmania, Australia |
| SRR1663553 | Ithaca, USA | SRR1663596 | Tasmania, Australia |
| SRR1663554 | Ithaca, USA | SRR1663597 | Tasmania, Australia |
| SRR1663555 | Ithaca, USA | SRR1663598 | Zimbabwe |
| SRR1663556 | Ithaca, USA | SRR1663599 | Zimbabwe |
| SRR1663557 | Ithaca, USA | SRR1663600 | Zimbabwe |
| SRR1663558 | Ithaca, USA | SRR1663601 | Zimbabwe |
| SRR1663559 | Ithaca, USA | SRR1663602 | Zimbabwe |
| SRR1663560 | Ithaca, USA | SRR1663603 | Zimbabwe |
| SRR1663561 | Netherlands | SRR1663604 | Zimbabwe |
| SRR1663562 | Netherlands | SRR1663605 | Zimbabwe |
| SRR1663563 | Netherlands | SRR1663606 | Zimbabwe |
| SRR1663564 | Netherlands | SRR1663607 | Zimbabwe |
| SRR1663565 | Netherlands | SRR1663608 | Zimbabwe |
| SRR1663566 | Netherlands | SRR1663609 | Zimbabwe |
| SRR1663567 | Netherlands | SRR1663610 | Zimbabwe |
| SRR1663568 | Netherlands | SRR1663611 | Zimbabwe |
| SRR1663569 | Netherlands | | |

Table 3: Overview of the most abundant diagnostic SNPs, for identifying population structure of Tirant, 412 and Opus. For each diagnostic SNP we show the position in the TE (position), the population where the SNP is most abundant (population), a frequency threshold and the abundance of the SNP inside (in) and outside (out) of the given population. For example, the first entry shows that the SNP in Tirant at position 5628 has a frequency of at least 0.5 (threshold) in 68% of the samples from Tasmania (in) but only in 10% of the other samples (i.e. not-Tasmania; out).

| **TE** | **position** | **population** | **threshold** | **in** | **out** |
|---|---|---|---|---|---|
| Tirant | 5628 | Ithaca, USA | 0.5 | 68% | 10% |
| Tirant | 6757 | Ithaca, USA | 0.5 | 63% | 19% |
| Tirant | 2254 | Ithaca, USA | 0.3 | 63% | 0% |
| Tirant | 243 | Tasmania, Australia | 0.5 | 94% | 20% |
| Tirant | 275 | Tasmania, Australia | 0.5 | 100% | 11% |
| Tirant | 3921 | Tasmania, Australia | 0.5 | 94% | 0% |
| Tirant | 4374 | Tasmania, Australia | 0.5 | 83% | 0% |
| Tirant | 8350 | Tasmania, Australia | 0.5 | 100% | 16% |
| Tirant | 8382 | Tasmania, Australia | 0.5 | 100% | 9% |
| Tirant | 5091 | Tasmania, Australia | 0.4 | 72% | 0% |
| Tirant | 3733 | Beijing, China | 0.4 | 60% | 7% |
| Tirant | 1547 | Beijing, China | 0.3 | 80% | 3% |
| Tirant | 4311 | Netherlands | 0.3 | 58% | 20% |
| 412 | 7199 | Zimbabwe | 0.5 | 79% | 4% |
| 412 | 7168 | Zimbabwe | 0.4 | 64% | 0% |
| 412 | 115 | Zimbabwe | 0.3 | 79% | 3% |
| 412 | 1530 | Zimbabwe | 0.3 | 71% | 6% |
| 412 | 1536 | Zimbabwe | 0.3 | 71% | 7% |
| 412 | 1542 | Zimbabwe | 0.3 | 71% | 7% |
| 412 | 1554 | Zimbabwe | 0.3 | 57% | 6% |
| 412 | 369 | Beijing, China | 0.4 | 53% | 11% |
| 412 | 5700 | Beijing, China | 0.4 | 60% | 11% |
| 412 | 7422 | Beijing, China | 0.4 | 60% | 13% |
| Opus | 325 | Tasmania, Australia | 0.5 | 78% | 6% |
| Opus | 7328 | Tasmania, Australia | 0.5 | 78% | 7% |
| Opus | 7369 | Zimbabwe | 0.5 | 71% | 0% |
| Opus | 366 | Zimbabwe | 0.4 | 79% | 0% |
| Opus | 6385 | Ithaca, USA | 0.4 | 84% | 20% |

Table 4: TE content in four long-read assemblies of *D. melanogaster*. For each of the seven TEs that invaded *D. melanogaster* populations during the last 200 years we show the genomic proportion in kbp. The assembly size (size) and the genomic proportion occupied by these seven TEs is also shown.

| | Pi2 | DGRP-732 | Iso1 | Canton-S |
|---|---|---|---|---|
| size | 167,834 | 141,551 | 143,726 | 149,105 |
| 412 | 312 | 234 | 242 | 249 |
| Blood | 260 | 99 | 218 | 261 |
| Opus | 263 | 167 | 248 | 221 |
| Tirant | 69 | 84 | 161 | 0 |
| I-element | 81 | 123 | 68 | 0 |
| Hobo | 188 | 55 | 33 | 0 |
| P-element | 58 | 41 | 0 | 0 |
| Sum TEs | 1,230 | 803 | 970 | 731 |
| proportion [%] | 0.73 | 0.57 | 0.68 | 0.49 |

Table 5: Overview of the long-read assemblies of diverse insect species analysed in this work.

| order | family | genus | taxon | accession |
|---|---|---|---|---|
| Coleoptera | Coccinellidae | Cryptolaemus | Cryptolaemus.montrouzieri | GCA_013387265.1 |
| Coleoptera | Coccinellidae | Harmonia | Harmonia.axyridis | GCA_011033045.1 |
| Coleoptera | Coccinellidae | Propylea | Propylea.japonica | GCA_013421045.1 |
| Coleoptera | Curculionidae | Listronotus | Listronotus.bonariensis | GCA_014170235.1 |
| Coleoptera | Curculionidae | Sitophilus | Sitophilus.oryzae | GCA_002938485.2 |
| Coleoptera | Elateridae | Limonius | Limonius.californicus | GCA_014611495.1 |
| Coleoptera | Lampyridae | Abscondita | Abscondita.terminalis | GCA_013368085.1 |
| Coleoptera | Lampyridae | Lamprigera | Lamprigera.yunnana | GCA_013368075.1 |
| Coleoptera | Lampyridae | Photinus | Photinus.pyralis | GCA_008802855.1 |
| Coleoptera | Nitidulidae | Aethina | Aethina.tumida | GCA_001937115.1 |
| Coleoptera | Scarabaeidae | Protaetia | Protaetia.brevitarsis | GCA_004143645.1 |
| Coleoptera | Scarabaeidae | Trypoxylus | Trypoxylus.dichotomus | GCA_014905495.1 |
| Coleoptera | Silphidae | Nicrophorus | Nicrophorus.vespilloides | GCA_001412225.1 |
| Collembola | Entomobryidae | Sinella | Sinella.curviseta | GCA_004115045.2 |
| Collembola | Isotomidae | Folsomia | Folsomia.candida | GCA_002217175.1 |
| Collembola | Orchesellidae | Orchesella | Orchesella.cincta | GCA_001718145.1 |
| Diptera | Calliphoridae | Cochliomyia | Cochliomyia.hominivorax | GCA_004302925.1 |
| Diptera | Calliphoridae | Phormia | Phormia.regina | GCA_001735545.1 |
| Diptera | Chironomidae | Belgica | Belgica.antarctica | GCA_000775305.1 |
| Diptera | Culicidae | Aedes | Aedes.albopictus | GCA_001876365.2 |
| Diptera | Culicidae | Aedes | Aedes.aegypti | GCA_002204515.1 |
| Diptera | Culicidae | Anopheles | Anopheles.albimanus | GCA_013758885.1 |
| Diptera | Culicidae | Anopheles | Anopheles.gambiae | GCA_001542645.1 |
| Diptera | Culicidae | Anopheles | Anopheles.funestus | GCA_003951495.1 |
| Diptera | Culicidae | Anopheles | Anopheles.stephensi | GCA_013141755.1 |
| Diptera | Culicidae | Anopheles | Anopheles.coluzzii | GCA_004136515.2 |
| Diptera | Diopsidae | Teleopsis | Teleopsis.dalmanni | GCA_002237135.1 |
| Diptera | Muscidae | Haematobia | Haematobia.irritans | GCA_003123925.1 |
| Diptera | Muscidae | Musca | Musca.domestica | GCA_014843735.1 |
| Diptera | Sarcophagidae | Sarcophaga | Sarcophaga.peregrina | GCA_014635995.1 |
| Diptera | Sciaridae | Bradysia | Bradysia.coprophila | GCA_014529535.1 |
| Diptera | Tephritidae | Bactrocera | Bactrocera.oleae | GCA_001188975.4 |
| Hemiptera | Aleyrodidae | Bemisia | Bemisia.tabaci | GCA_001854935.1 |
| Hemiptera | Aleyrodidae | Trialeurodes | Trialeurodes.vaporariorum | GCA_011764245.1 |
| Hemiptera | Anthocoridae | Orius | Orius.insidiosus | GCA_014119065.1 |
| Hemiptera | Aphididae | Aphis | Aphis.glycines | GCA_009928515.1 |
| Hemiptera | Aphididae | Rhopalosiphum | Rhopalosiphum.maidis | GCA_003676215.3 |
| Hemiptera | Aphididae | Sitobion | Sitobion.miscanthi | GCA_008086715.1 |
| Hemiptera | Delphacidae | Laodelphax | Laodelphax.striatellus | GCA_003335185.2 |
| Hemiptera | Delphacidae | Nilaparvata | Nilaparvata.lugens | GCA_014356525.1 |
| Hemiptera | Liviidae | Diaphorina | Diaphorina.citri | GCA_000475195.1 |
| Hemiptera | Miridae | Apolygus | Apolygus.lucorum | GCA_009739505.2 |
| Hemiptera | Pentatomidae | Euschistus | Euschistus.heros | GCA_003667255.1 |
| Hemiptera | Pseudococcidae | Maconellicoccus | Maconellicoccus.hirsutus | GCA_003261595.1 |
| Hemiptera | Pseudococcidae | Phenacoccus | Phenacoccus.solenopsis | GCA_009761765.1 |
| Hemiptera | Reduviidae | Triatoma | Triatoma.infestans | GCA_011037195.1 |
| Hymenoptera | Apidae | Apis | Apis.mellifera | GCA_003254395.2 |
| Hymenoptera | Apidae | Apis | Apis.dorsata | GCA_009792835.1 |
| Hymenoptera | Apidae | Apis | Apis.mellifera | GCA_003314205.1 |
| Hymenoptera | Apidae | Apis | Apis.cerana | GCA_011100585.1 |
| Hymenoptera | Apidae | Apis | Apis.mellifera | GCA_013841205.1 |
| Hymenoptera | Apidae | Apis | Apis.mellifera | GCA_013841245.1 |
| Hymenoptera | Apidae | Bombus | Bombus.vosnesenskii | GCA_011952255.1 |
| Hymenoptera | Apidae | Bombus | Bombus.bifarius | GCA_011952205.1 |
| Hymenoptera | Apidae | Bombus | Bombus.vancouverensis | GCA_011952275.1 |
| Hymenoptera | Braconidae | Aphidius | Aphidius.gifuensis | GCA_014905175.1 |
| Hymenoptera | Braconidae | Aphidius | Aphidius.ervi | GCA_011426455.1 |
| Hymenoptera | Braconidae | Chelonus | Chelonus.insularis | GCA_013357705.1 |
| Hymenoptera | Braconidae | Lysiphlebus | Lysiphlebus.fabarum | GCA_011426435.1 |
| Hymenoptera | Colletidae | Colletes | Colletes.gigas | GCA_013123115.1 |
| Hymenoptera | Figitidae | Leptopilina | Leptopilina.clavipes | GCA_001855655.1 |
| Hymenoptera | Figitidae | Leptopilina | Leptopilina.boulardi | GCA_011634795.1 |
| Hymenoptera | Formicidae | Camponotus | Camponotus.floridanus | GCA_003227725.1 |
| Hymenoptera | Formicidae | Formica | Formica.selysi | GCA_009859135.1 |
| Hymenoptera | Formicidae | Harpegnathos | Harpegnathos.saltator | GCA_003227715.1 |
| Hymenoptera | Formicidae | Monomorium | Monomorium.pharaonis | GCA_013373865.2 |
| Hymenoptera | Formicidae | Nylanderia | Nylanderia.fulva | GCA_005281655.1 |
| Hymenoptera | Formicidae | Ooceraea | Ooceraea.biroi | GCA_003672135.1 |
| Hymenoptera | Formicidae | Solenopsis | Solenopsis.invicta | GCA_010367695.1 |
| Hymenoptera | Megachilidae | Osmia | Osmia.lignaria | GCA_012274295.1 |
| Hymenoptera | Pteromalidae | Nasonia | Nasonia.vitripennis | GCA_009193385.2 |
| Hymenoptera | Pteromalidae | Pteromalus | Pteromalus.puparum | GCA_012977825.2 |
| Hymenoptera | Vespidae | Polistes | Polistes.metricus | GCA_010416925.1 |
| Hymenoptera | Vespidae | Polistes | Polistes.fuscatus | GCA_010416935.1 |
| Hymenoptera | Vespidae | Vespa | Vespa.mandarinia | GCA_014083535.1 |
| Lepidoptera | Carposinidae | Carposina | Carposina.sasakii | GCA_014607495.2 |
| Lepidoptera | Crambidae | Chilo | Chilo.suppressalis | GCA_004000445.1 |
| Lepidoptera | Crambidae | Cnaphalocrocis | Cnaphalocrocis.medinalis | GCA_014851415.1 |
| Lepidoptera | Hesperiidae | Epargyreus | Epargyreus.clarus | GCA_014595695.1 |
| Lepidoptera | Lasiocampidae | Dendrolimus | Dendrolimus.punctatus | GCA_012273795.1 |
| Lepidoptera | Noctuidae | Heliothis | Heliothis.virescens | GCA_002382865.1 |
| Lepidoptera | Noctuidae | Spodoptera | Spodoptera.exigua | GCA_011316535.1 |
| Lepidoptera | Noctuidae | Spodoptera | Spodoptera.frugiperda | GCA_011064685.1 |
| Lepidoptera | Noctuidae | Trichoplusia | Trichoplusia.ni | GCA_003590095.1 |
| Lepidoptera | Nymphalidae | Maniola | Maniola.jurtina | GCA_009667785.1 |
| Lepidoptera | Pieridae | Colias | Colias.croceus | GCA_009982905.1 |
| Lepidoptera | Psychidae | Eumeta | Eumeta.japonica | GCA_005406025.1 |
| Lepidoptera | Pyralidae | Galleria | Galleria.mellonella | GCA_004355975.1 |
| Lepidoptera | Saturniidae | Antheraea | Antheraea.mylitta | GCA_014332785.1 |
| Lepidoptera | Saturniidae | Samia | Samia.ricini | GCA_014132275.1 |
| Lepidoptera | Sphingidae | Hyles | Hyles.vespertilio | GCA_009982885.1 |
| Lepidoptera | Sphingidae | Manduca | Manduca.sexta | GCA_014839805.1 |
| Lepidoptera | Tortricidae | Cydia | Cydia.pomonella | GCA_003425675.2 |
| Orthoptera | Gryllidae | Teleogryllus | Teleogryllus.occipitalis | GCA_011170035.1 |
| Siphonaptera | Pulicidae | Ctenocephalides | Ctenocephalides.felis | GCA_003426905.1 |
| Thysanoptera | Thripidae | Thrips | Thrips.palmi | GCA_012932325.1 |
| Trichoptera | Hydropsychidae | Hydropsyche | Hydropsyche.tenuis | GCA_009617725.1 |
| Trichoptera | Polycentropodidae | Plectrocnemia | Plectrocnemia.conspersa | GCA_009617715.1 |
| Trichoptera | Stenopsychidae | Stenopsyche | Stenopsyche.tienmushanensis | GCA_008973525.1 |

### References

S. Luo, H. Zhang, Y. Duan, X. Yao, A. G. Clark, and J. Lu. The evolutionary arms race between transposable elements and pirnas in drosophila melanogaster. *BMC Evolutionary Biology*, 20(1):1–18, 2020.

F. Schwarz, F. Wierzbicki, K.-A. Senti, and R. Kofler. Tirant stealthily invaded natural drosophila melanogaster populations during the last century. *bioRxiv*, 2020.

L. Weilguny and R. Kofler. DeviaTE: Assembly-free analysis and visualization of mobile genetic element composition. *Molecular ecology resources*, 19(5):1346–1354, 2019.
